# Microbial dispersal from surrounding vegetation influences phyllosphere microbiome assembly of corn and soybean

**DOI:** 10.1101/2025.05.31.657201

**Authors:** Kyle M. Meyer, Steven E. Lindow

## Abstract

Non-crop plants surrounding large plantings of agricultural crops can provide numerous ecological services to adjacent agricultural plants but have rarely been considered as a source of microorganisms during the early stages of their growth. In this study we test whether agricultural plants in close proximity to surrounding woodland habitat fragments develop a denser microbiome than plants farther away, and whether the composition of the crop microbiomes more closely resembles the composition of the surrounding vegetation when in close proximity. During the early stages of development, we sampled epiphytic bacteria from corn and soybean leaves over 4 and 3 weekly sampling timepoints, respectively, using a spatially explicit design, and on the final timepoint for both host species we additionally sampled a younger cohort of leaves. To contextualize the source strength of the surrounding vegetation we also sampled the soil at each sampling location. Both crop species exhibited a microbiome density gradient and a decay of microbiome similarity to surrounding vegetation over a distance of 100 m from the field edges at many timepoints. Phyllosphere microbiome similarity to the soil also tended to increase into the field interior. The strength of host plant microbiome filtering also depended on the proximity to the surrounding vegetation, with intermediate to most distant locations exhibiting the highest values of host filtering, reflecting an apparent decrease in immigrant inoculum. The bacterial communities of younger leaves tended to more closely resemble those of the older surrounding conspecific leaves than either the soil or surrounding woodland vegetation, reflecting the growing dominance of inoculum from within developing crop canopies as plants grew. Overall, our study sheds light on the important role that dispersal of bacteria from nearby leaves can play in phyllosphere microbiome assembly and highlights the diminishing role that soil plays in assembly of phyllosphere microbiomes as plant sources are closer or more abundant.

## Introduction

Communities of bacteria, fungi, and viruses residing on leaf surfaces, termed the phyllosphere microbiome (Lindow and Brandl, 2003), form *de novo* on newly emerged tissues via the arrival of microbial propagules from the environment and subsequent host filtering. Host filtering takes place by means of plant immune activity, molecular signaling, and/or the development of species-specific chemical and physical features (Bodenhausen et al., 2014; Custer et al., 2022; Hacquard et al., 2017; Horton et al., 2014; Humphrey and Whiteman, 2020; Jones and Dangl, 2006; Yadav et al., 2005; Zhalnina et al., 2018), and these characteristics can give rise to predictable differences in microbiome composition among hosts (Bodenhausen et al., 2014; Meyer et al., 2023; Morella et al., 2020; Wagner et al., 2019, 2016), a phenomenon termed “species identity effects”. The net result of host filtering tends to be an enrichment of microorganisms with traits adapted to high UV exposure and temperature extremes, as well as water and nutrient resource scarcity (Vorholt, 2012). The phyllosphere microbiome is known to have direct effects on the health outcome of the plant, including disease resistance, by promoting competitive exclusion of pathogens and/or the maturation of plant innate immune systems, and these protective mechanisms often depend on the recruitment of key specialized taxa (Innerebner et al., 2011; Lindow et al., 2024; Morella et al., 2019; Ritpitakphong et al., 2016; Vogel et al., 2016, 2021). Phyllosphere microbiomes can also improve plant reproductive success (Mehlferber et al., 2023). Thus, the formation and functionality of the phyllosphere microbiome, including its host benefits, may depend on the suitability of microbial colonists as well as the timing of their arrival from the surrounding sources.

A major challenge in the study of phyllosphere microbiology is identifying the sources from which the microbiome members have emigrated. A commonly purported source of phyllosphere microorganisms is the soil, and this is supported by observations that phyllosphere microbiomes often share high taxonomic overlap with soil microbiomes (Tkacz et al., 2020; Zarraonaindia et al., 2015) and that certain microorganisms are capable of migrating from soil to the phyllosphere, particularly to leaves in close proximity to the soil (Massoni et al., 2021).

Other studies, however, have shown that the influence of soil taxa on the phyllosphere decreases over the growing season (Copeland et al., 2015), or is less important relative to epiphytic communities on nearby vegetation (Whitaker et al., 2023, 2021). Indeed, fluxes of microorganisms from vegetation have been reported in air parcels above forested, rural, and urban areas (Docherty et al., 2018; Galès et al., 2014; Lymperopoulou et al., 2016; Mhuireach et al., 2016; Womack et al., 2015), and the identity and biomass of a plant’s nearby neighbors have been shown to impact the diversity, composition and host specialization of its phyllosphere (Lajoie and Kembel, 2021; Meyer et al., 2023, 2022). Lastly, in a greenhouse setting, stochastic effects such as sources of airborne inoculum earlier in plant colonization have been shown to strongly influence microbiome assembly on leaves (Maignien et al., 2014).

One might expect the surrounding vegetation to be a more influential source of microorganisms than the soil, since it would more likely be enriched in phyllosphere-adapted taxa, but this may depend on the state of the environment into which the plant emerges. For instance, in large-scale agricultural practices monocultures are typically planted all at once and all other plants within such large fields are suppressed by herbicides or cultivation; thus, as seedlings emerge there is little standing vegetation to act as a source of plant-associated microbial propagules. Phyllosphere microbiomes of seedlings raised in monoculture might therefore tend to be more influenced by the soil unless they emerge near a patch of established vegetation. Incorporating greenspace such as woodlands, prairie, or hedgerows into agroecosystems has been shown to provide numerous ecological services to the crops and the surrounding environment, including boosting pollinator and natural enemy populations (Amy et al., 2015; Garratt et al., 2017; Staley et al., 2023). What is less clear, however, is whether such local patches of vegetation can act as a substantial source of phyllosphere microorganisms.

Here we test three main hypotheses: 1) established vegetation surrounding annual crop fields acts as a strong source of microorganisms during the early stages of crop phyllosphere microbiome assembly, 2) the magnitude of the influence of soil on phyllosphere microbiome assembly depends on the developing plant’s proximity to vegetation and the host’s age/developmental stage, and 3) the microbial communities on young leaves emerging into a more developed population of conspecific plants will be more influenced by such plants than by soil or nearby woodland vegetation, due to presumed enrichment of host-specialized bacteria. We test these hypotheses using corn and soybean plants in an experimental agricultural setting in Rosemount, Minnesota, USA over 4 and 3 weekly sampling timepoints, respectively, with a spatially explicit sampling scheme in which sampling locations were designated at 0, 30, and 100 m from surrounding mixed woodland vegetation in replicate sites. Our results shed light on the spatial dynamics of phyllosphere microorganisms in an agroecological context and provide spatial and temporal perspectives on the relationship between the similarities of microbiomes of soil and those on the leaves of developing crop plants and that on surrounding vegetation.

## Methods

### Experimental Design

To test the hypothesis that vegetation surrounding agricultural sites can act as an important source of microorganisms during early phyllosphere microbiome assembly, we performed an observational field study involving corn (*Zea maize*), soybean (*Glycine max*), and composite leaf material samples from surrounding woodlands at the University of Minnesota Rosemount Research and Outreach Center in Rosemount, Minnesota between 31 May and 30 June, 2022 (Fig. 1A). Within this research and outreach center, we selected 5 sites where corn was planted, and 5 where soybean was planted into large (> 30 ha) fields that were surrounded by large areas (> 50 ha in extent) of a mixture of short herbaceous vegetation and moderate to tall woody vegetation. The emergence of plants other than either of the 2 crop plant species was completely blocked in all of the fields by pre-plant herbicides. We established two sampling transects separated by approximately 30 m at each site. Each transect had 3 sampling locations: one starting at the edge of the fields adjacent to the source vegetation (0 m), one 30 m into the field in the presumed direction of the prevailing wind, and one 100 m into the field along the same path. Composite samples of source vegetation were collected adjacent to the 0 m sampling location of each transect and at one location in between the transects, thus giving a total of 9 samples per site at each sampling time: 6 crop phyllosphere samples, and 3 composite source vegetation phyllosphere samples. Composite samples of the surrounding vegetation included leaves from box elder (*Acer negundo*), black ash (*Fraxinus nigra*), American basswood (*Tilia americana*), black walnut (*Juglans nigra*), silver maple (*Acer saccharinum*), white oak (*Quercus alba*), ironwood (*Ostrya virginiana*), pin cherry (*Prunus pensylvanica*), rock elm (*Ulmus thomasii*), buckthorn (*Rhamnus cathartica*), and various grasses.

**Fig. 1:**
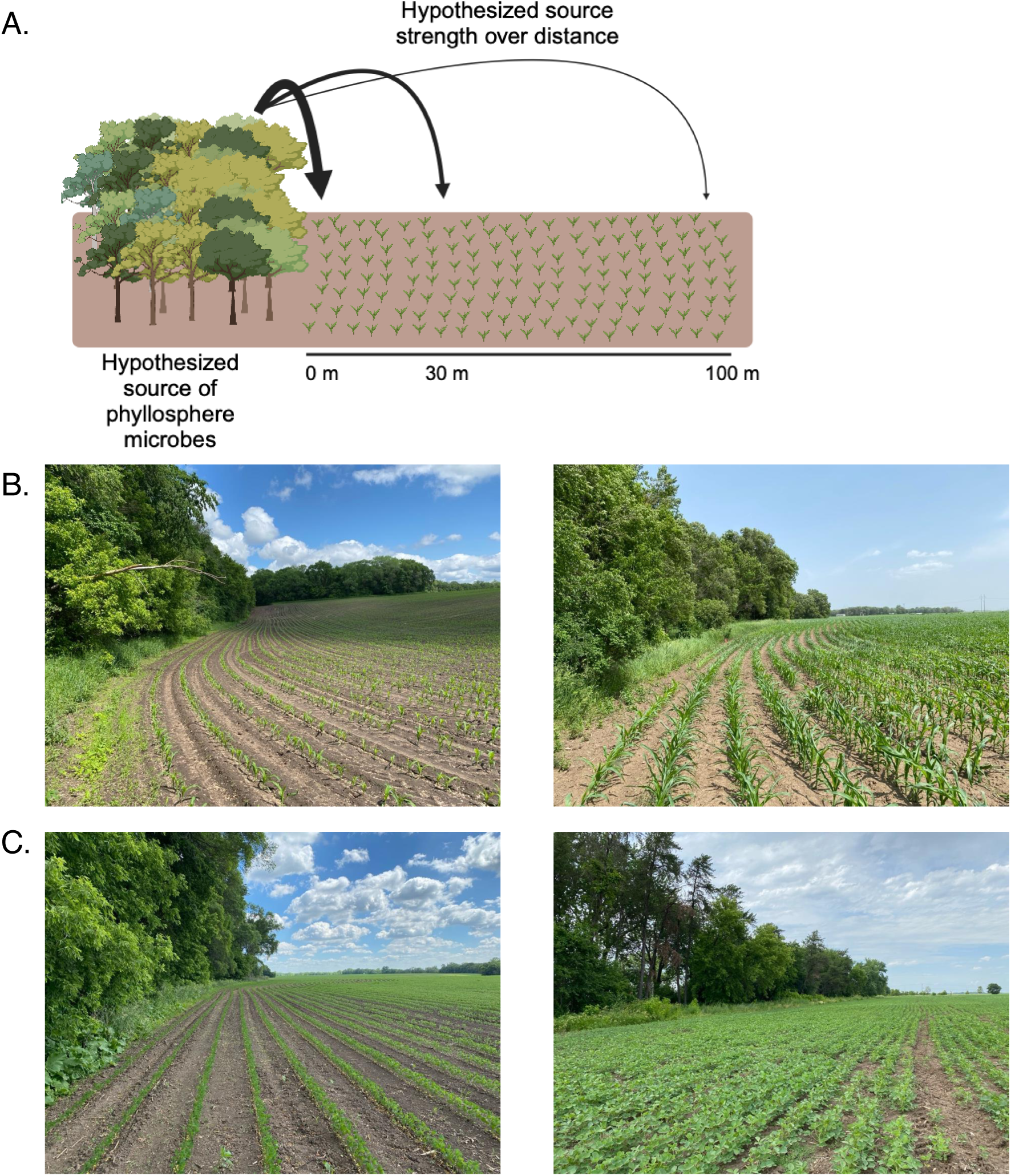
Conceptual outline of the experiment. A) Hypothesis of the experiment: woodland vegetation will act as a source of leaf-adapted microbial propagules to the nearby crops (corn and soybean), and the strength of this source with decay over space, giving a distance-decay pattern of phyllosphere microbial densities and compositional similarity to that of the vegetation. B) Photographs of corn plants at timepoint 1 (left) and timepoint 4 (right). C) Photographs of soybean plants at timepoint 1 (left) and timepoint 3 (right). Panel A was created using Biorender.

Corn was sampled weekly on 4 occasions (see Fig. 1B for site photos), and soybean was sampled weekly on 3 occasions (Table 1, see Fig. 1C for site photos). For the first sampling of both crops, soil samples were collected from each sampling location to test the contribution of soil microbial taxa to phyllosphere microbiome composition. To survey the soil, we used an ethanol-sterilized trowel to sample the top 2 cm of soil at three areas within a sampling location. These three samples from each sampling location were combined into a composite sample in a sterile Ziplock bag and placed on ice immediately in the field and then subsequently frozen the same night.

**Table 1:**
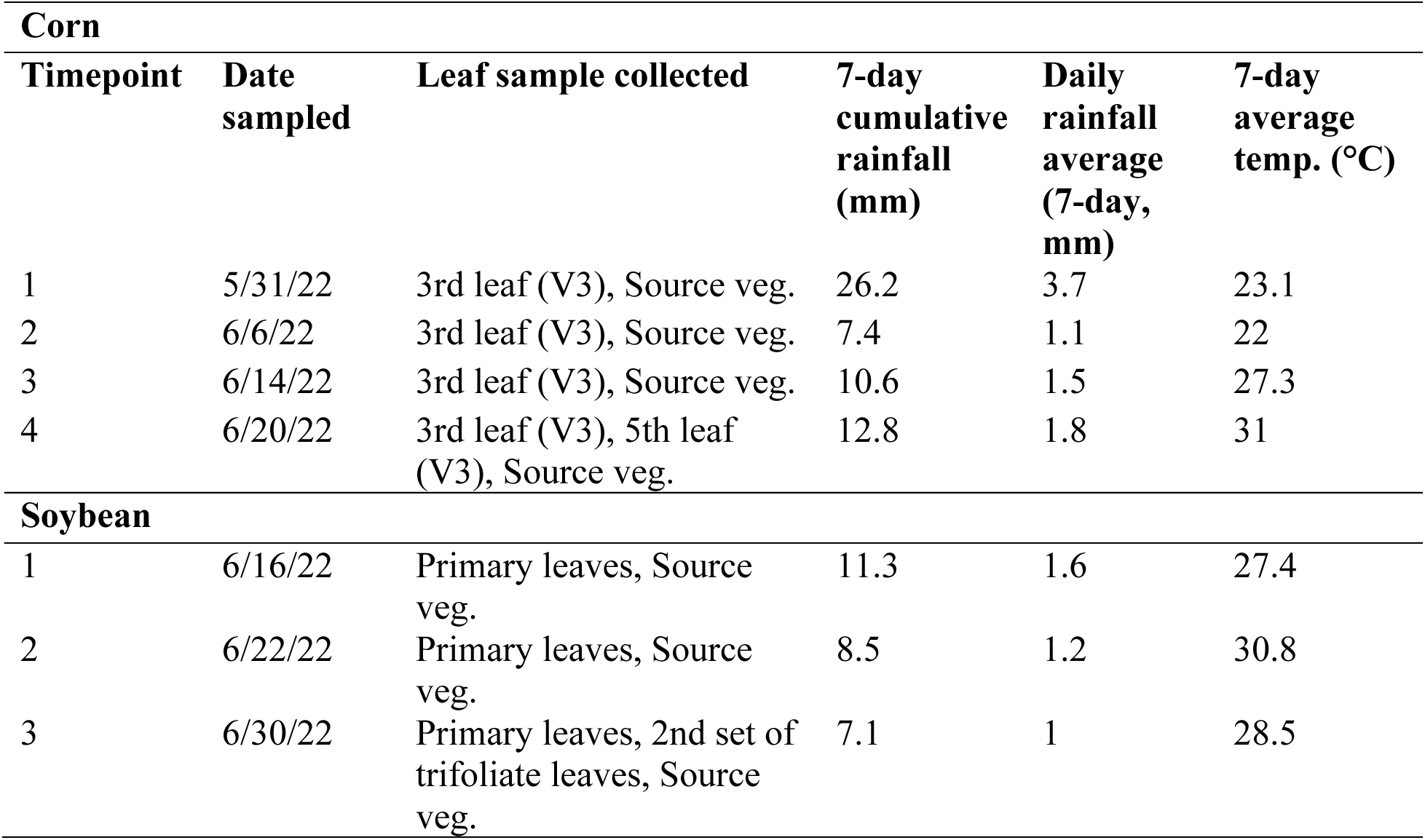
Summary of leaf collection campaigns for corn and soybean samples. Meteorological data (precipitation and temperature) pertain to the day of sampling and the 6 preceding days.

Corn plants were in the V3 developmental stage at the first sampling period and had reached the V7 developmental stage by the last sampling period. At each sampling location, the third leaf was collected from approximately 10 adjacent plants, forming a composite sample.

Composite samples of this leaf cohort were collected at each time point from plants adjacent to those that had been sampled previously. In this way we consistently surveyed leaves from the same developmental stage as the host plants became older. At time point 4, we separately collected both the third and fifth leaves from the same cohort of plants to compare the effects of leaf age on phyllosphere microbiome composition. The soybean plants were at the V1 developmental stage (emergence of first trifolate leaves) and had reached the R3 developmental stage by the last sampling period. Composite samples of the primary leaves were collected at each sampling location following the same procedure as the corn. At time point 3, we separately collected the primary and second set of trifolate leaves to compare the effects of leaf age on phyllosphere microbiome composition.

Leaves of both crop species and surrounding vegetation were removed from the plants using ethanol-sterilized nitrile gloves, placed in sterile Ziplock bags, and kept in a chilled cooler in the field. Leaf samples were kept at 4° C overnight before processing the following morning in a laboratory at the University of Minnesota. Epiphytic microbial communities were gently dissociated from the leaf surfaces by adding 10 mM MgCl2 to each sample (60 ml for crop species, 75 ml for composite vegetation) and sonicating in either a Branson 2510 sonicating water bath or a Cole-Parmer Ultrasonic Cleaner for 10 minutes. The resultant leaf wash was concentrated by centrifuging at 3220 rcf and 4° C for 20 minutes, followed by decanting the supernatant. The pellet was then resuspended in 2 ml sterile 10 mM MgCl2, aliquoted into two

### 2.5 ml tubes, and frozen for subsequent DNA extraction and bacterial cell enumeration

#### DNA extraction, library preparation, and sequencing

To survey the abundance and diversity of the bacterial phyllosphere, we extracted DNA from each of the leaf wash samples. Half of the frozen resuspended leaf wash was used for DNA extraction using the DNeasy Powersoil Pro kit (Qiagen, Germany) following manufacturer’s instructions. Sample order was randomized to avoid batch effects, and a blank (no sample) control was included in each round of DNA extraction. DNA concentration was quantified using a Qubit dsDNA HS assay kit (Thermo Scientific, USA). Library preparation and sequencing were performed by Novogene. Briefly, sample DNA was used as template and PCR amplified for 30 cycles using the 799F (5’ – AACMGGATTAGATACCCKG – 3’) - 1193R (5’ – ACGTCATCCCCACCTTCC – 3’) primer combination, which targets the V5-V7 region of the 16S rRNA gene and minimizes chloroplast amplification. PCR reactions were carried out with 15 μL of Phusion® High -Fidelity PCR Master Mix (New England Biolabs), 2 μM of forward and reverse primers, and approximately 10 ng sample DNA. Thermal cycling consisted of initial denaturation at 98° C for 1 min, followed by 30 cycles of denaturation at 98° C for 10 s, annealing at 50° C for 30 s, and elongation at 72° C for 30 s, and 72° C for 5 min. PCR products were then run on a 1% agarose gel and purified using the Qiagen Gel Extraction Kit (Qiagen, Germany).

Sequencing libraries were constructed with index codes using the TruSeq DNA PCR-Free Sample Preparation Kit (Illumina, USA) following manufacturer’s recommendations. The library quality was assessed on a Qubit 2.0 Fluorometer (Thermo Scientific, USA) and an Agilent Bioanalyzer 2100 system. Lastly, the library was sequenced on the Illumina NovaSeq platform with 250 bp paired-end reads.

#### Sequence processing

The resultant 16S rRNA gene amplicon sequences were processed using the DADA2 pipeline implemented in the R statistical environment, including the packages ShortRead, Biostrings, and Phyloseq. Forward and reverse reads were truncated at 220 and 220 bp, respectively, and quality filtered using the function ‘filterAndTrim’ with default settings (i.e. maxN=0, maxEE=c(2,2)), and truncQ=2). Error rates for forward and reverse reads were determined using the ‘learnErrors’ function, and then applied to remove sequencing errors from reads and assign them to amplicon sequence variants (ASVs) using the ‘dada’ function. Paired reads were merged, converted into a sequence table, and then chimeric sequences were removed from the sequence table. Taxonomy was assigned to the remaining ASVs using the ‘assignTaxonomy’ function, which implements the RDP Naïve Bayesian Classifier algorithm with kmer size 8 and 100 bootstrap replicates. This taxonomic classification used the Silva (version 138) SSU taxonomic training dataset formatted for DADA2. Chloroplast and mitochondrial sequences were filtered from the ASV table by removing any ASVs with a taxonomic assignment of ‘Chloroplast’ at the Order level or ‘Mitochondria’ at the Family level, respectively. Lastly, we applied the ‘isContaminant’ function (method = prevalence) from the package ‘decontam’ to our samples using the blank (no sample) DNA extractions to identify and remove putative contaminants introduced during DNA extraction.

#### Bacterial cell enumeration using droplet digital PCR

Foliar bacterial abundances of plant samples were estimated using droplet digital PCR (ddPCR) on the Bio-Rad QX200 system (Bio-Rad, USA). We targeted the V5-V7 region of the 16S rRNA gene of cells in the leaf washings using the same 799 F–1193R primer combination as the amplicon sequencing described above. Five microlitres of 1:10 diluted leaf wash was combined with 11 μl of 2× EvaGreen Supermix (Bio-Rad, USA) and 0.22 μl of each primer, and 5.56 μl of molecular grade water to a total volume of 20 μl. Reaction mixes were then loaded into the QX200 droplet generator with 70 μl of droplet generation oil and then transferred to a PCR plate. Thirty-nine cycles of PCR were performed under the following conditions: 95 °C for 10 min, 95 °C for 30 s, 55°C for 30 s, 72 °C for 2 min, with steps 2–4 repeated 39 times, 4 °C for 5 min, and 90 °C for 5 min. EvaGreen signal was measured on the QX200 droplet reader, cutoff thresholds were set for each column based on background fluorescence in no template controls, and concentrations were determined using the associated QuantaSoft software.

### Statistical analysis

All statistical analyses were performed using R version 4.3.2 (R Core Team, 2020). Community matrices were rarefied to 23500 counts per sample ten times and averaged to account for differences in sampling extent across samples. In the case where soil-affiliated taxa were removed (Fig. 4), communities were rarefied to 1340 counts per sample. ASV rarefaction curves were generated using the ‘rarecurve’ function in the vegan package in R (Oksanen et al., 2015). Bray Curtis bacterial community dissimilarities were calculated between samples using the ‘vegdist’ function, also in the vegan package. To test compositional differences among soybean and corn phyllosphere microbiomes we used a PERMANOVA, implemented by the ‘adonis2’ function in the vegan package and adjusted the *p* values for subsetting using the ‘p.adjust’ function (method = ‘hochberg’). The relationship between bacterial abundance and geographic distance from the surrounding vegetation was assessed using a linear mixed effect model, with distance as the fixed effect and site as a random effect, using the ‘lmerTest’ package (Kuznetsova et al., 2017). The same procedure was used to assess the distance-decay of similarity to the surrounding vegetation. Univariate data such as bacterial cells per g leaf material and community similarity to sources were analyzed using an anova and Tukey’s HSD post-hoc analysis, after confirming the normal distribution of model fit residuals. The differential abundance of bacterial taxa at the 0m sampling locations relative to the 100m sampling locations was performed using DESeq2 (Love et al., 2014). All figures were produced using ggplot2 (Wickham, 2009).

## Results

### Study overview

The dataset contained 423 samples, with 31,021,784 total sequencing reads after quality filtering and removal of chimeras, resulting in a total of 308,340 amplicon sequence variants (ASVs), 67,635 of which were detected more than 5 times, and 32,841 of which were detected more than 10 times. The corn phyllosphere dataset had the highest number of taxa detected (101,891 ASVs, 19,327 were detected more than 10 times), followed by soybean (81,522 ASVs, 12,818 were detected more than 10 times), soil (60,350 ASVs, 13,370 were detected more than 10 times), and then the composite surrounding vegetation samples (46,740 ASVs, 10,989 were detected more than 10 times). Supplementary Fig. 1A, B, and C show compositional summaries at the order, family, and genus levels for the surrounding vegetation, soil, corn, and soybean samples. In all four sample types, the rarefaction curves level-off, indicating that the sequencing efforts have achieved adequate sampling depth (Supp. Fig. 2).

### Phyllosphere microbiome density decreases with distance from vegetation at certain timepoints

To test the hypothesis that proximity to source vegetation influences the density of a plant’s microbiome, we estimated epiphytic bacterial cell density of each leaf wash sample using droplet digital PCR (ddPCR). Cell counts per g leaf material were significantly higher on the surrounding vegetation (8.6 x 10^6^ ± 1.3 x 10^6^) than on the corn (5.0 x 10^5^ ± 1.1 x 10^5^) or the soybean (1.9 x 10^5^ ± 6.9 x 10^4^) (Fig. 2A, Supp. Fig. 3, Tukey’s HSD, *p* < 0.001 for both). In the soybean samples, epiphytic microbiome densities were consistently lower than on the surrounding source vegetation at all three distances sampled (*p* < 0.001, for all three distances). The same can be said for the corn samples, where epiphytic microbiome densities at all three sampling locations were lower in density than on the surrounding vegetation (source – 0 m, *p* < 0.05; source – 30 m, *p* < 0.01; source – 100 m, *p* < 0.01).

**Fig. 2:**
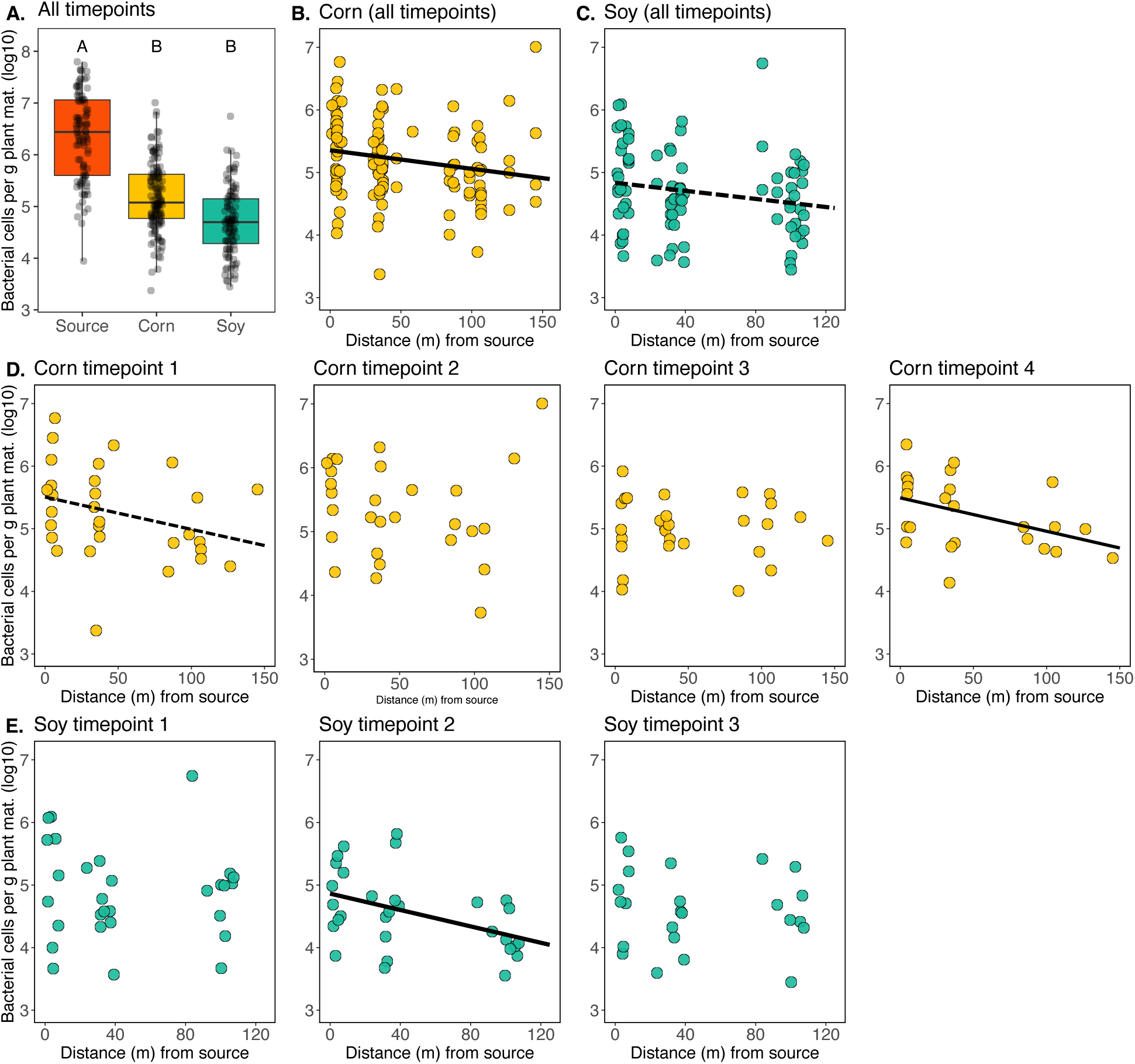
F**a**ctors **influencing densities of epiphytic microbiome on corn and soybean.** A) Epiphytic bacterial cells per g plant material (y-axis) differ among host species (x-axis), with “source” denoting leaves from the surrounding woodland vegetation. Significant differences (letters above boxes) based on Tukey’s HSD. The relationship between epiphytic bacterial cells per g leaf material (y-axis) and distance from the surrounding woodland vegetation (x-axis) for corn (B) and soybean (C) samples, with all time points combined. The solid line indicates a statistically significant relationship (*p* < 0.05), while the dashed line indicates a statistically trending relationship (*p* < 0.1). D, E) The relationship between epiphytic bacterial cells per g plant material (y-axis) and distance from the surrounding woodland vegetation (x-axis) broken down by time point for corn (D) and soybean (E). Significant relationships (Corn timepoint 4 & Soybean timepoint 2, *p* ≤ 0.05) are denoted by a solid fit line, while a statistically trending relationship (Corn timepoint 1, *p* = 0.09) is denoted by a dashed fit line.

When the corn dataset across the four timepoints are combined a negative linear relationship with distance from the source vegetation is seen (Fig. 2B, R^2^ = 0.03, *p* < 0.05). A similar relationship in the combined soybean data is seen, but with less statistical support (Fig. 2C, R^2^ = 0.03, *p* = 0.08). This same relationship is seen at certain individual time points. For instance, in the corn samples at timepoint 4 and the soybean samples at timepoint 2, distance from surrounding vegetation is a significant predictor of microbiome density (Fig. 2D,E: corn timepoint 4: R^2^ = 0.15, *p* < 0.05, soybean timepoint 2: R^2^ = 0.16, *p* < 0.01). In other words, at these time points, the closer a plant was to the surrounding vegetation, the denser its phyllosphere microbiome was. For the corn samples at timepoint 1, a similar trend is seen, but with less statistical significance (R^2^ = 0.06, *p* = 0.08).

To put these linear relationships into perspective, for the corn samples at timepoint 1, given a linear fit of y = -0.0052x + 5.51, a five-fold decrease in bacterial cell density, i.e. from log10(5.5) to log10(5.0), occurred over a 96.8 m distance from the source vegetation. Likewise for the corn samples at timepoint 4, given a linear fit of y = -0.0053x + 5.49, a five-fold decrease in bacterial cell density occurred over 94.3 m of distance from the source vegetation. Finally, for the soybean samples at timepoint 2, given a linear fit of y = -0.0065x + 4.86, a five-fold decrease in bacterial cell density occurred over 76.4 m.

Finally, we asked whether plants directly adjacent to the surrounding vegetation developed epiphytic microbiomes more quickly over the timepoints sampled, relative to those farther into the field. This was not the case for either host plant, where bacterial cell densities per g of leaf were indistinguishable at the 0 m sampling locations across the timepoints sampled (*p* > 0.05, for both cases).

### The relationship between phyllosphere and soil microbiomes depends, in part, on distance from surrounding vegetation

To better understand the strength of the soil microbiome as a source of phyllosphere microorganisms, we next examined pairwise similarity between each phyllosphere sample and the soil sample corresponding to its sampling location. From this analysis, overall high levels of similarity between soil and phyllosphere membership was seen (Corn: range 0.05 - 0.60, median = 0.42, Soy: range 0.23 - 0.61, median = 0.48). For the corn samples multiple linear regression revealed a positive relationship between the distance from surrounding vegetation and the similarity of the phyllosphere microbiome to the soil microbiome (Fig. 3A, t = -1.95, *p* = 0.05). In other words, the farther the samples were into the field interior, the more closely their microbiomes resembled the soil. A hump-shaped relationship was seen with sampling timepoint (Fig 3B), whereby similarity to the soil microbiome increased from time point 1 to 2 (t = -4.93, *p* < 0.001), returned to initial values at timepoint 3 (t = 3.83, *p* > 0.05), and then further decreased at timepoint 4 (*p* < 0.001), with no effect of site or any interactions therein. For the soybean samples, a similar positive relationship was seen between a sample’s distance from the surrounding vegetation and its similarity to the soil microbiome, but with less statistical support (Fig. 3C, t = -1.79, *p* = 0.07), as well as an interaction between distance from surrounding vegetation and site (Fig.3D), where samples at site S2 exhibited an opposite spatial trend (t = 2.60, *p* < 0.05).

**Fig. 3:**
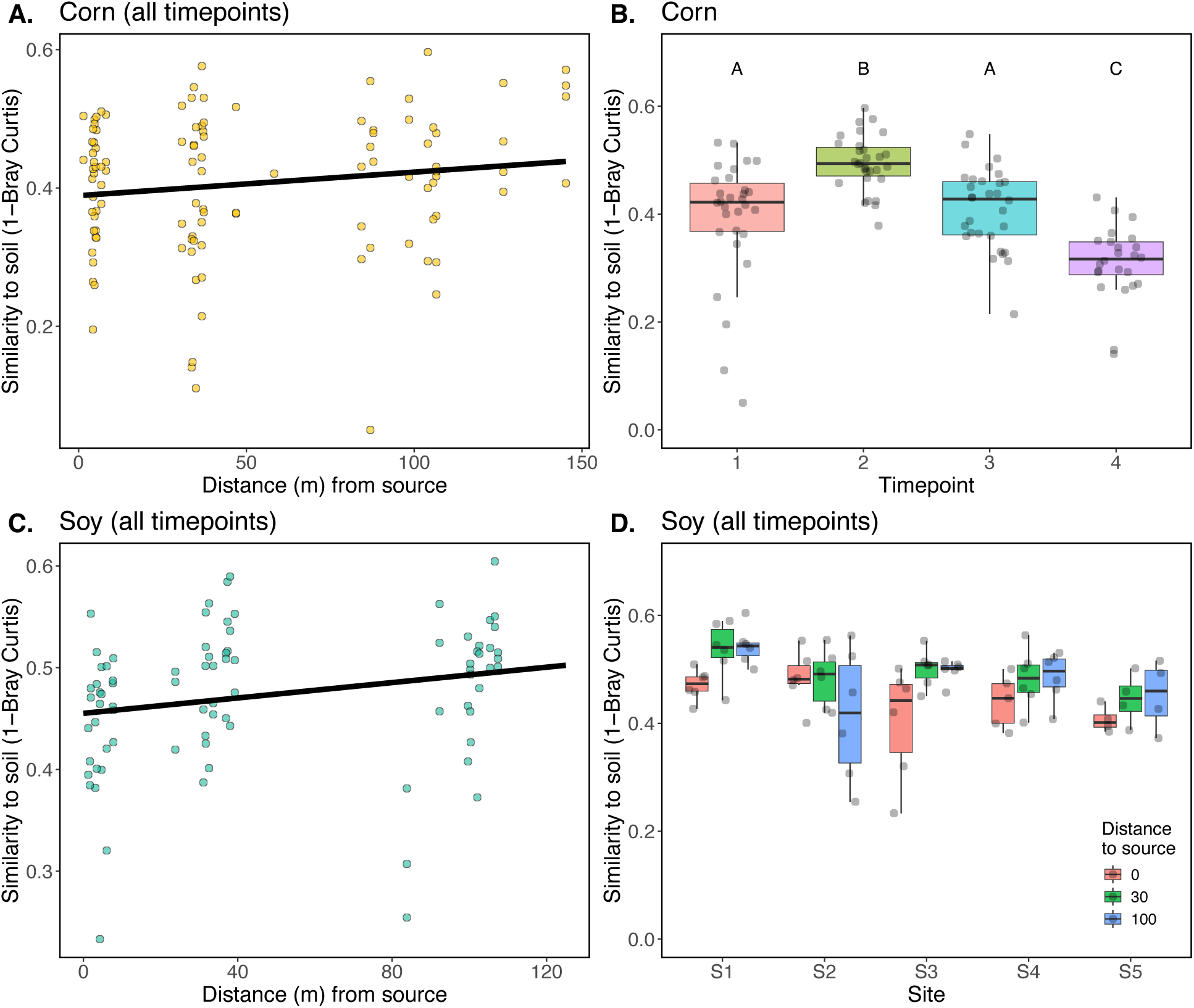
T**h**e **relationship between phyllosphere and soil bacterial microbiomes depends on the host plant’s distance from the surrounding vegetation.** A) Corn phyllosphere microbiome similarity (1-Bray Curtis, y-axis) to soil increases with distance (x-axis) from the surrounding woodland vegetation. All timepoints are shown. B) Corn phyllosphere similarity to soil (y-axis) follows a hump-shaped pattern through time (x-axis), letters represent Tukey’s HSD significant differences. C) Soybean phyllosphere microbiome similarity (1-Bray Curtis, y-axis) to soil increases with distance (x-axis) from the vegetation surrounding fields. All timepoints are shown. D) Soybean phyllosphere similarity to soil (y-axis) exhibits a distance to surrounding vegetation (box color) x site (x-axis) interaction.

**Fig. 4:**
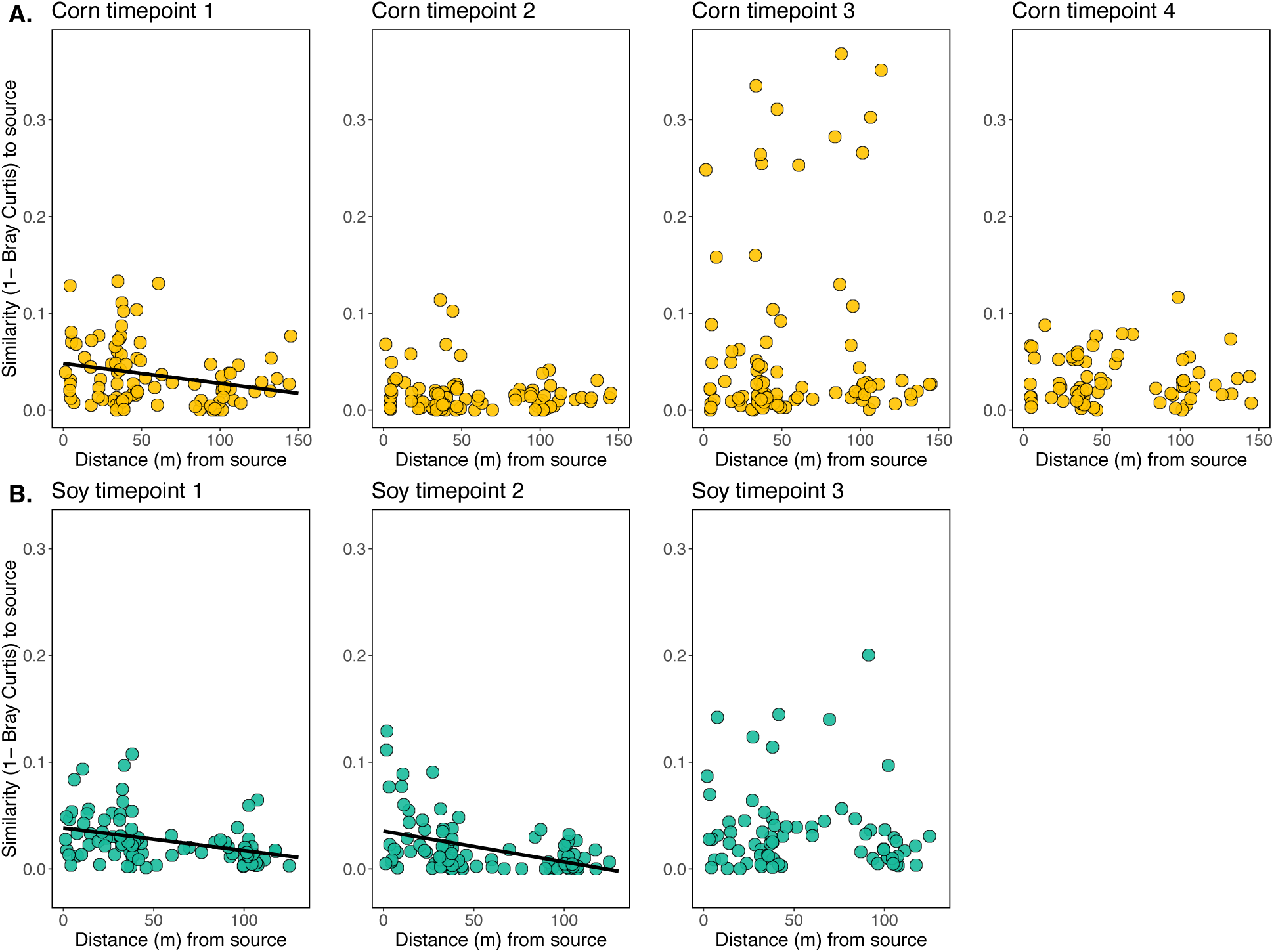
Relationship between epiphytic microbiome similarity to surrounding source vegetation (1-Bray Curtis, y-axis) and distance (m) from source vegetation (x-axis), after removing taxa detected in corresponding soil samples. Solid fit lines are shown where a statistically significant (*p* < 0.05) relationship is observed. See Supp. Fig. 4 for the same comparisons, but where soil taxa were not removed.

We hypothesized that many of the soil taxa detected in the phyllosphere had passively dispersed onto the nearby plant surfaces following heavy rains in the days preceding sampling, but may not be forming established, multiplying populations. We therefore performed differential abundance analysis to identify the subset of soil taxa that were enriched, thus representing a higher relative abundance, in the phyllosphere relative to the soil. This approach identified 135 taxa that were significantly enriched in the soybean samples (Supp. Table 1), and 212 taxa that were significantly enriched in the corn samples (Supp. Table 1). In both cases, most of these taxa were either present in a very low relative abundance in the soil and/or detected in very few of the soil samples. Furthermore, hypothesizing that phyllosphere microbiomes laden with soil microbes would grow to lower densities, we asked whether there was a relationship between phyllosphere microbiome similarity to soil and its bacterial cell density. This, however, was not the case for corn (*p* > 0.05) or soybean (*p* > 0.05) samples.

### Proximity to source vegetation ephemerally influences crop phyllosphere bacterial microbiome composition

We next tested whether a crop’s proximity to the surrounding vegetation influenced its phyllosphere microbiome composition. We hypothesized that the crop phyllosphere microbiome compositions would exhibit a distance-decay pattern of similarity to the source vegetation adjacent to fields. Soybean phyllosphere bacterial microbiome structure exhibited a statistically significant distance-decay pattern of community similarity to the surrounding vegetation at timepoints 1 and 2 (Supp. Fig. 4A,B, *p* < 0.05 and *p* = 0.001, respectively). In other words, soybean plants that were closer to surrounding vegetation early in their development tended to be more compositionally similar to the microbial communities found in that vegetation, than plants that were farther into the fields. This pattern was no longer detectable at timepoint 3 (*p* > 0.05). Corn phyllosphere microbiome structure did not exhibit the hypothesized distance-decay pattern at timepoints 1 (*p* > 0.05), 2 (*p* > 0.05), or 3 (*p* > 0.05); and the relationship was statistically trending at timepoint 4 (*p* = 0.08).

We hypothesized that soil particles on the leaves, and thus soil microbiome members, could be masking the distance-decay signal in the phyllosphere microbiomes, so we removed the taxa identified in our soil samples and again performed distance-decay analysis. This process removed an average of 89.9 ± 5.2 % of the soybean bacterial communities, 86.7 ± 9.2 % of the corn bacterial communities, and 84.4 ± 15.6% of the surrounding vegetation bacterial communities. For the corn phyllosphere samples, this procedure revealed a significant distance-decay pattern at timepoint 1 (Fig. 4A, *p* < 0.001). For the soybean phyllosphere samples, we see no qualitative change to our conclusion of distance decay after removing taxa detected in the soil, but we see stronger statistical support for such a relationship at timepoints 1 and 2 (Fig. 4B, *p* = 0.001 and *p* = 0.002, respectively). Thus, it appears that soilborne bacteria found on the leaf surfaces may in part mask some of the microbiome similarity to the surrounding vegetation.

We additionally asked whether the taxa detected in the surrounding vegetation exhibited a decreasing density gradient into the field interior. We first calculated the collective relative abundance of the vegetation-associated bacteria in each sample and then multiplied this value by each sample’s corresponding ddPCR-inferred bacterial abundance (assessed using the same primer combination). The estimated absolute abundances of these vegetation-associated taxa on corn leaf surfaces ranged from 1.3 x 10^3^ to 5.3 x 10^6^ cells per g leaf material (median = 7.8 x 10^4^) and exhibited a decreasing density abundance with distance into the field interior at timepoint 4 (*p* < 0.05, Supp. Fig 5A) and at timepoint 1, but with less statistical support (*p* = 0.07, Supp. Fig 5A). For the soybean leaf samples, epiphytic densities of vegetation-affiliated bacteria ranged from 2.0 x 10^3^ to 3.7 x 10^6^ cells per g leaf material (median = 3.3 x 10^4^) and exhibited a density gradient into the field interior at timepoint 2 (*p* = 0.01, Supp. Fig 5B). For the corn samples, this density gradient is also apparent when all four timepoints are combined (*p* = 0.02, Supp. Fig 5C), but this is not the case for the soybean samples (*p* > 0.05, Supp. Fig 5D).

### Microbiome compositional differences between corn and soybean strongest at intermediate or farthest distance from surrounding vegetation

We next hypothesized that corn and soybean plants closer to the woodland edge would have more opportunity for ecological filtering due to the increased prevalence of leaf-associated microbial propagules, and hence would exhibit stronger host species identity effects, as exemplified by larger host effects (R^2^ values) in a PERMANOVA model. We subsetted corn and soybean samples to match their time of sampling and their distance from surrounding vegetation and then compared their compositions using a PERMANOVA. At timepoints 1 and 2 we see a hump-shaped curve of host species identity effects whereby the corn and soybean samples at locations at the edge of fields near surrounding vegetation are the least distinguishable, while the those sampled 30 m from the edge are the most distinguishable, and those located 100 m from the edge were intermediate in effect (Fig. 5, orange and green lines, all *Padj* < 0.05). At timepoint 3 we see a larger and monotonic increase in species identity effects with distance from the surrounding vegetation (Fig. 5, purple line, all *Padj* < 0.05). Thus, at all sampling times corn and soybeans nearest surrounding vegetation exhibited the weakest species identity effects, and samples at the intermediate or most interior locations exhibited the strongest.

**Fig. 5:**
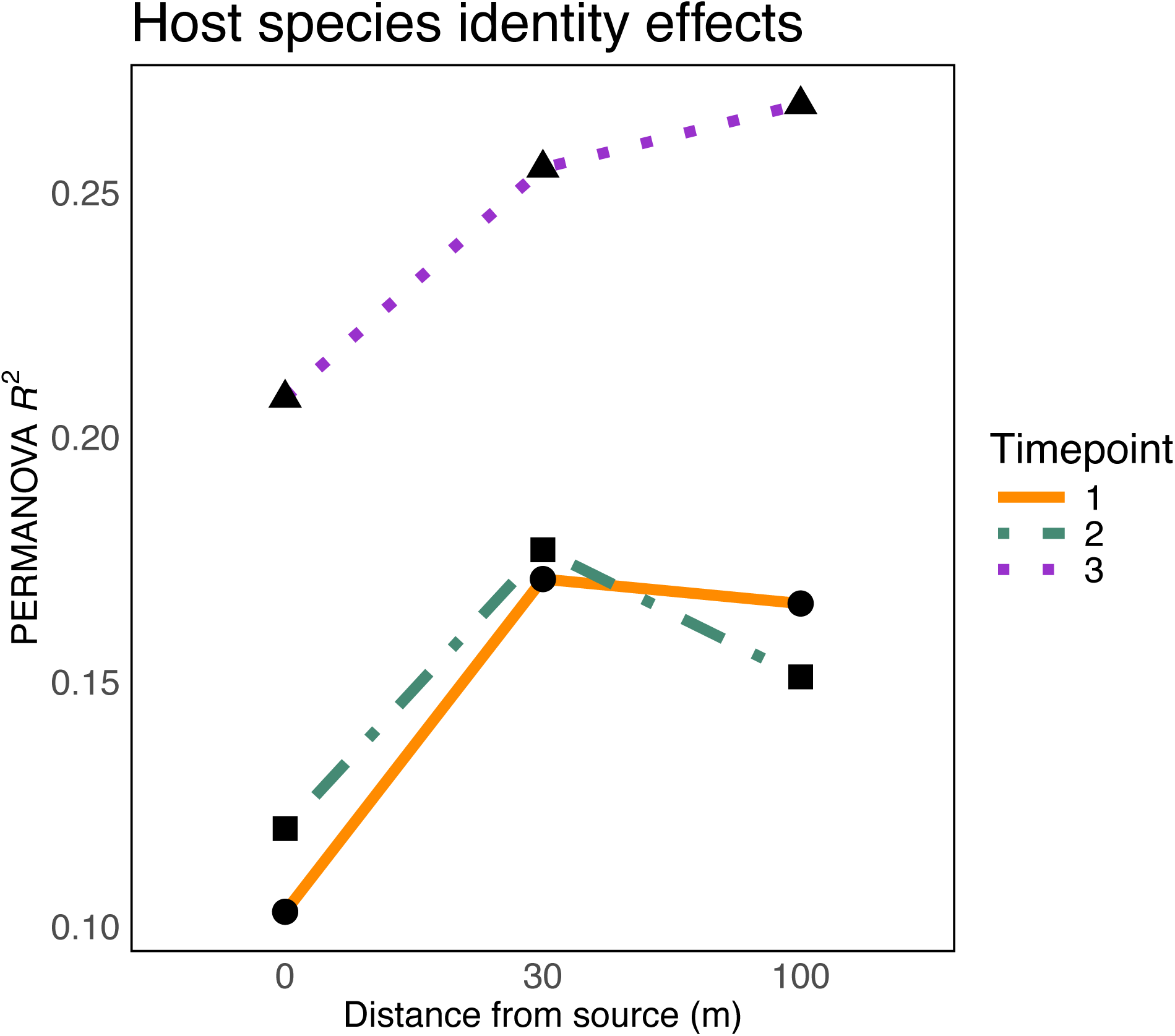
Compositional differences (PERMANOVA R^2^ values, y-axis) between corn and soybean bacterial phyllosphere microbiomes at timepoints 1 (orange solid line, circle points), 2 (green dot-dash line, square points), and 3 (purple dotted line, triangle points) separated by their distance (m) from the vegetation surrounding fields (x-axis). All R^2^ values presented are statistically significant (*p*adj < 0.05).

### Phyllosphere microbiome homogeneity increases towards field interior

We next hypothesized that the crop plants farthest into the interior of the field would have the most homogeneous phyllosphere microbiomes, due to the uniformity of the surrounding plant community. To test this, we examined Bray Curtis dissimilarity levels within microbiomes at the 0, 30, and 100 m sampling locations for each sampling time and crop species. We find support for our hypothesis for both crops at some, but not all, timepoints. For instance, corn microbiomes were more homogeneous at the 100 m sampling locations than the 0 m at timepoints 3 and 4 (Tukey’s HSD *p* < 0.001 & *p* < 0.05, respectively, Supp. Fig. 6A). Additionally, at timepoints 1 and 3 the soybean microbiomes at the 100 m sampling locations were significantly more homogenous than those immediately adjacent to surrounding vegetation (Tukey’s HSD *p* < 0.001, Supp. Fig. 6B). At timepoint 2, only the 30 m soybean microbiomes were more homogeneous than those immediately adjacent to surrounding vegetation (Tukey’s HSD *p* = 0.01). It thus appears that at a majority of sampling times, the plants farther into the interior of the field, which are thus surrounded by conspecific plants, tend to develop more homogeneous phyllosphere microbiomes.

### Taxa that are differentially abundant near vegetation are detected in vegetation, but also soil

We next asked whether plants at the 0 m sampling locations were enriched in taxa from the source vegetation, relative to those 100 m interior to the fields at those sample times at which distance-decay relationships with the vegetation were detected. For the soybean samples at timepoint 1, we identified 12 taxa belonging to 3 bacterial classes (Gammaproteobacteria, Actinobacteria, and Thermoleophilia, see Supp. Table 2 for finer taxonomic affiliations), that were differentially enriched in the 0 m samples relative to the 100 m samples. Of these 12, 11 were detected in the source vegetation samples and 10 were also detected in the soil samples. For the soybean plants sampled at timepoint 2, we identified 13 taxa that were more abundant in the 0 m versus the 100 m samples. These were affiliated with the bacterial classes Gammaproteobacteria and Actinobacteria (Supp. Table 2). Of these 13 taxa, 12 were detected in the source vegetation, and all 13 were also detected in the soil. For the corn samples at timepoint 1, we identified 8 taxa that were differentially enriched in the 0 m locations relative to the 100 m locations. These taxa were affiliated with the classes Gammaproteobacteria, Actinobacteria, and Myxococcia (Table 2). All 8 taxa were also detected in both the vegetation and soil samples.

### Young leaves at later timepoints are more similar to older leaves than to the soil or surrounding vegetation

At the last sampling time for each crop species, when the crop canopies had become more developed, we also collected a younger cohort of leaves at each sampling location in addition to the regular older, leaves to ask whether the microbial communities on these more recently emerged leaves might be more influenced by the local microbial communities on the older corn/soybean leaves and hence be less influenced by the vegetation surrounding the field. We further hypothesized that since these younger leaves were emerging farther from the soil surface, the resultant leaf microbiomes would be less influenced by the soil microbial community.

We first assessed the strength of three putative sources of microorganisms emigrating onto the leaves of the young leaves: older leaves of the same crop, nearby soil, and nearby vegetation. To do so, we calculated the pairwise compositional similarity (1- Bray Curtis) between each sample and the older leaves from the same sampling location and time, the soil from the same sampling location, and the vegetation surrounding the croplands at the same time and transect as the sample. For both corn and soybean samples, we observed a statistical difference between microbiome similarities to the three putative sources of microorganisms (Corn: F2,69 = 98.1, *p* < 0.001, Soy: F2,69 = 408.8, *p* < 0.001). For both crops we see the same pattern: the microbiomes of young leaves are the most compositionally similar to their older leaf counterparts (Tukey’s HSD *p* < 0.001), followed by soil (Tukey’s HSD *p* < 0.001), followed lastly by surrounding vegetation (Tukey’s HSD *p* < 0.001) (Fig. 6A,B). For comparison, we assessed the soil and surrounding vegetation as sources of microbial propagules for the older leaves from these time points, revealing that for both corn and soybean the soil is the stronger source than the surrounding vegetation (F1,52 = 98.4, *p* < 0.001, F1,46 = 282.9, *p* < 0.001, Supp. Fig. 7A,B).

**Fig. 6:**
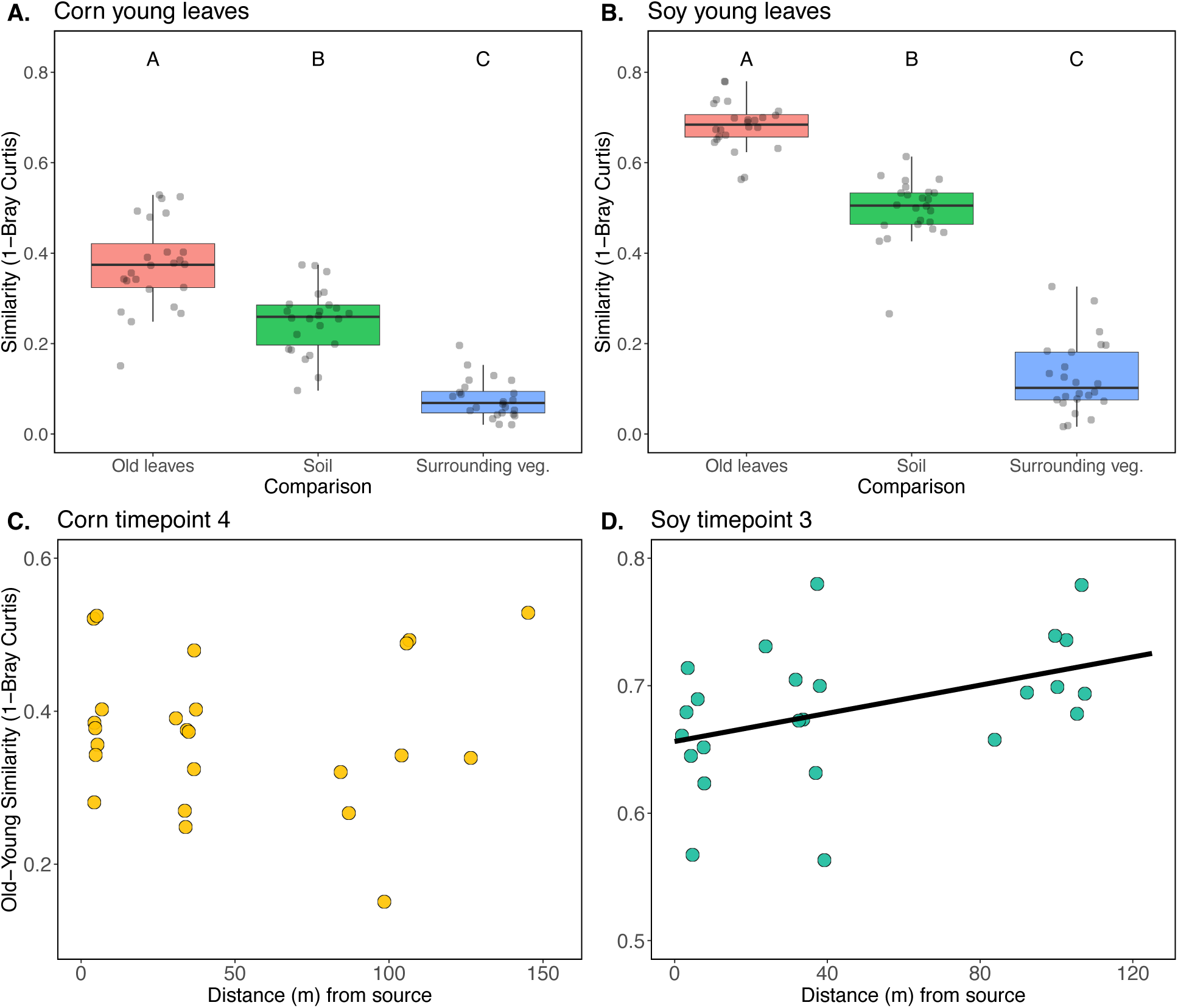
Taxonomic similarity (1-Bray Curtis, y-axis) of epiphytic microbiomes from young corn (A) and soybean (B) to three putative sources of microbial propagules (x-axis): older leaves from the same sampling location and time (Old leaves), soil from the paired sampling locations (Soil), and leaves from the vegetation surrounding the field sampled at the same transect and timepoint (Surrounding veg.). C) Pairwise similarity (1-Bray Curtis, y-axis) between old and young corn and soybean (D) leaves sampled from the same cohort of leaves at a given location. Distance from source(x-axis) indicates how far away the sampling location was from the vegetation at the edge of each field.

### Similarities of old and young soybean leaf microbiomes increase with distance from vegetation surrounding fields

We next hypothesized that a host plant’s proximity to the surrounding vegetation would impact the similarity between the old and young leaves. More specifically, we hypothesized that young leaves from plants father into the field interior would be more influenced by the surrounding older crop leaves, whereas young leaves closer to the surrounding vegetation would be colonized by a wider assortment of microbial propagules from both the surrounding vegetation and the surrounding crops, resulting in less similarity between the old and young leaves at a given sampling location. We find support for this hypothesis in the soybean samples (*p* < 0.05), but not the corn samples (Fig. 6C,D). In other words, soybean leaf bacterial communities on plants further into the field interior have higher taxonomic similarity to those from the older crop leaves that surround them than to those located near the edge of fields with other plant species nearby.

### Young soybean leaf phyllospheres more closely resemble contemporaneous older leaves than microbiomes at earlier timepoints

Next, we asked which microbiome that had developed over time on a particular plant species most resembled that found on young leaves, to gain insights into whether the communities on young leaves more closely resembled those from earlier in plant growth or whether they reflected later successional taxa. To do so, we compared the young leaf microbiomes to those on leaves from plants sampled earlier in their growth at a given location in the field, as well as to the older leaves that were contemporaneously sampled. Young corn leaf microbiomes were equally similar to that on more basal (older) leaves sampled at any of the 4 different times (Fig. 7A, *p* > 0.05). By contrast, the microbiomes on young soybean leaves were most similar to that on contemporaneous older leaves sampled at timepoint 3 (Fig. 7B, Tukey’s HSD *p* < 0.001), followed by those sampled at timepoint 2 (Tukey’s HSD *p* < 0.001), and finally at timepoint 1 (Tukey’s HSD *p* < 0.001). Thus, it appears that the microbial communities on younger soybean leaves are more similar to that on established older leaves sampled at the same time and location than they were to the older cohort of leaves sampled earlier in the season.

**Fig. 7:**
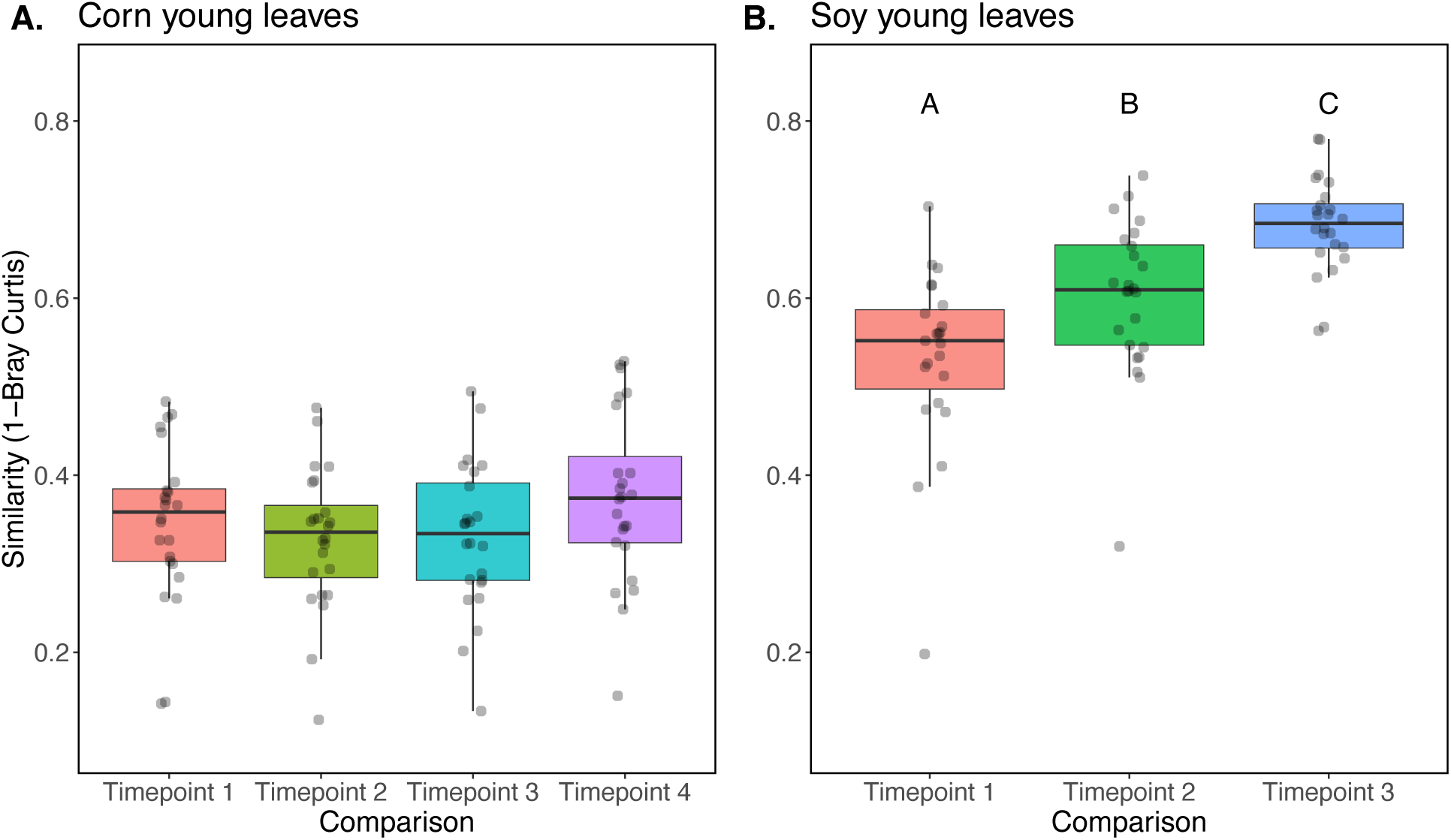
Taxonomic similarity (1-Bray Curtis, y-axis) between epiphytic microbiomes of young leaves from corn (A) and soybean (B) compared to that of an older cohort of leaves sampled previously in the season (x-axis) or at the same time (right most box in each plot).

### Distance-decay of microbiome similarity to surrounding vegetation exhibited in corn leaves following removal of soil taxa

We next asked whether the young leaf microbiomes exhibited a distance-decay pattern of similarity to the surrounding vegetation, similar to what was observed for the older leaves. For both the corn and soybean, these young leaf cohorts did not exhibit a statistically significant distance-decay pattern of microbiome similarity to the surrounding vegetation (*p* > 0.05). Similar to that seen on the older leaves, we hypothesized that ubiquitous soil microbiome members may be masking a decrease in similarity of epiphytic microbes with distance from putative sources on vegetation surrounding the fields. We thus, removed those taxa that were detected in soil samples and re-ran the analysis. This procedure revealed a statistically significant distance-decay pattern in the young corn leaves (Supp. Fig. 8A, p < 0.05), but not in the young soybean leaves (Supp Fig. 8B, *p* > 0.05).

**Fig. 8:**
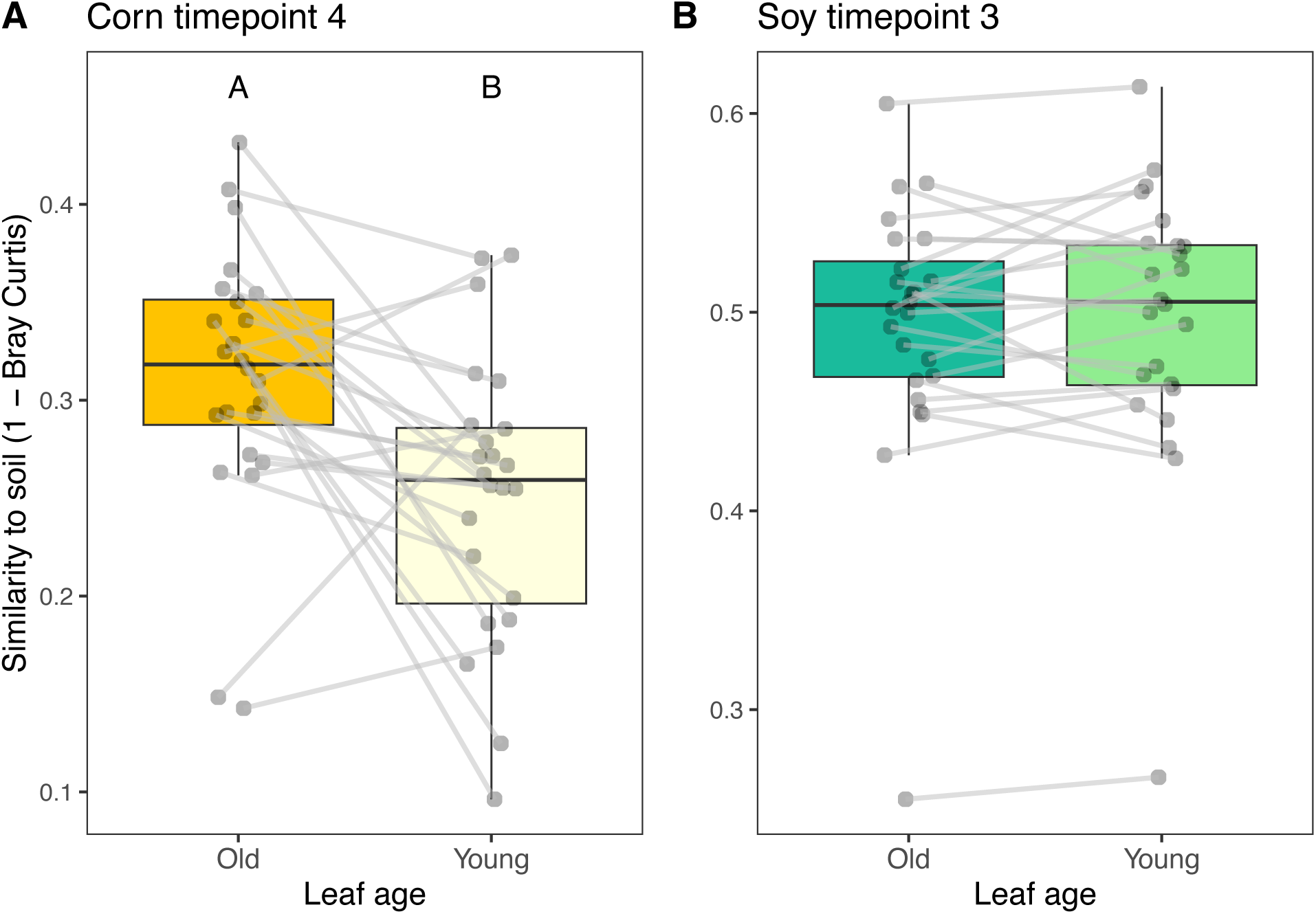
The taxonomic similarity (1-Bray, y-axis) between microbiomes on old and young corn (A) and soybean (B) leaves and the soil microbiomes at the same sampling locations. Lines connect samples from the same location. Letters indicate statistically significant differences (*p* < 0.05).

### Young corn leaf phyllosphere less similar to soil microbiome than older corn leaves

We next asked whether the bacterial communities on younger leaves would exhibit less compositional similarity to the soil microbial community relative to the older leaf cohorts, since the younger leaves tend to be farther from the soil surface. To do so, we compared the young and old leaf microbiomes to the soil microbiome from the corresponding sampling locations. This procedure revealed that the older leaves were more compositionally similar to soil than the younger leaves (*F*1,46 = 9.17, *p* < 0.01), (Fig. 8A). By contrast, the young and old soybean leaf cohorts did not differ in their similarity to the soil microbial community (*p* > 0.05, Fig. 8B).

### Young leaf phyllosphere bacterial cell densities indistinguishable from older leaves and are not impacted by proximity to woodland edge

Finally, we asked whether the younger leaves had lower epiphytic bacterial abundances relative to the older leaves, and whether proximity to vegetation surrounding the fields impacted such abundances. Cell densities per gram of plant material did not differ between young and old leaves of either corn or soybean (*p* > 0.05, Supp. Fig. 9A,B). Distance from surrounding vegetation was also not a significant predictor of epiphytic bacterial densities on young corn or soybean leaves.

## Discussion

In the present study, we asked whether established vegetation surrounding newly planted crop fields can act as an influential source of microbial colonists during the early stages of phyllosphere microbiome assembly on corn and soybean. We observed that the surrounding vegetation, which emerged at least 3 weeks prior to the crops, had 10 to100-fold greater bacterial densities than the crop plants (Fig. 2A), and that at certain sampling times for each crop (1 and 4 for corn, and 2 for soybean) a microbiome density gradient could be observed whereby epiphytic bacterial densities decreased by roughly 5-fold over our 100 m transects into the field interior (Fig. 2C,D). In addition to microbiome densities, we find evidence for neighboring vegetation effects on microbiome composition. For soybeans at timepoints 1 and 2, we see that plants close to the woodland edge share more microbiome similarities with that vegetation than those farther into the field interior (Supp. Fig. 4, Fig. 4B). In other words, we see a distance-decay of community similarity to woodland microbial communities. Of note, if we bioinformatically remove the taxa detected in soil from the leaf samples, then we also see evidence of this distance-decay relationship in corn (Fig. 4A, timepoint 1), suggesting that the woodland effect may be in part masked by the presence of soil particles (and hence soil taxa) on the leaf. We thus see that for two facets of microbial community structure (density and composition) a gradient of influence of neighboring vegetation is observed.

A growing body of work supports our observation that neighboring vegetation acts as an important source of microbial propagules to newly emerging plants or leaves. Lindow and Andersen (1996) showed that population sizes of ice nucleating bacteria were 6- to 30-fold higher on the leaves of orange trees that were in close proximity to nearby heterospecific vegetation, as opposed to other citrus trees. Meyer et al. (2022) showed that both the species identity and biomass of a plant’s immediate neighbors (<1 m away) can give rise to consistent distinctions in the focal plant’s foliar bacterial microbiome. Similarly, it has been shown that the presence of natural vegetative cover surrounding croplands has been associated with higher diversity of foliar fungi on crops (Whitaker et al., 2023), and that nearby plants are stronger sources of foliar fungi than soil (Whitaker et al., 2021). Our study contributes to this narrative in at least two ways: 1) it establishes that the abundance and composition of both corn and soybean phyllosphere microbiota can be influenced by surrounding vegetation at a scale of 0 to ∼100 m, and 2) it establishes that young leaves emerging on more developed plants are most influenced by the surrounding leaves of the same plant, or conspecific neighbors, than the soil or vegetation nearby fields (Fig. 6A,B). Other findings suggest that the transmission of foliar bacteria among conspecific plants can foster a competitive advantage for those taxa (i.e. host specialization) over taxa from a heterospecific host (Meyer et al., 2023). Thus, the predominant colonization of the young leaves by bacterial communities from the surrounding conspecific plants may be the result of an enrichment of host-specialized taxa that would have a competitive advantage over taxa from soil or heterospecific hosts.

By calculating species identity effects (*i.e.* microbiome composition distinctions among host species) we were able to examine the relationship between microbial dispersal from the vegetation nearby fields and host microbiome filtering on the crop leaf surfaces. Host filtering acts on the standing diversity of microbial taxa, therefore higher diversity and/or an adequate rate of immigration should provide sufficient opportunity for filtering to drive distinctions among host species. Metacommunity theory, however, predicts that under high rates of dispersal, optimal outcomes of host filtering may break down if taxa going locally extinct are continually replenished through dispersal (Leibold et al., 2022, 2004). High rates of dispersal are also associated with biotic homogenization, whereby communities become dominated by a common set of widespread taxa (Evans et al., 2017). Meyer et al. (2022) demonstrated empirical support for this theory by showing that the species identity effects of tomato, pepper, and common bean phyllosphere microbiomes were gradually diminished to nothing as their heterospecific neighbors increased in size and presumably exerted more propagule pressure on them. Meyer et al. (2023) additionally showed that repeated microbiome transmission among heterospecific hosts eroded species identity effects, likely due to the loss of host-specialized taxa and the proliferation of generalist taxa. Our present results contribute a spatial perspective to this narrative. For instance, at all three sampling times we see that host species identity effects are lower in plants closest to surrounding vegetation relative to the intermediate and farthest sampling locations. Our interpretation of this result is that high propagule pressure from the vegetation homogenized the two crop phyllospheres at the edge of the surrounding vegetation, but at locations farther into the field interior (30 m for timepoints 1 and 2, 100 m for timepoint 3, Fig. 5) host filtering may have benefitted from intermediate levels of dispersal, such that a sufficient supply of microorganisms could still arrive without driving microbiome homogenization. Beyond the observed spatial effect, we also see a temporal effect, such that species identity effects increased over time, likely due to more dispersal opportunities strengthening the outcome of host filtering and/or the maturation of host filtering mechanisms.

Similar changes to identity effects have been observed in the rhizosphere among sorghum cultivars over the growing season (Schlemper et al., 2017) and among grapevine rootstock microbiomes across host age difference (Berlanas et al., 2019). Thus, our results contribute to a growing narrative suggesting that species identity effects are subject to the influence of dispersal and host age.

Phyllosphere bacterial microbiomes have often been shown to exhibit high levels of similarity to soil microbiomes (Bodenhausen et al., 2013; Copeland et al., 2015; Massoni et al., 2021; Tkacz et al., 2020; Zarraonaindia et al., 2015). This high level of similarity could stem from at least three non-exclusive scenarios: 1) soil microbiomes contain taxa that can opportunistically colonize (and multiply on) leaves, 2) leaf microbiomes contain non-multiplying soil taxa that persist, or have died, but remain detectable through sequencing, and/or 3) the soil underneath a plant has become enriched in leaf-associated bacteria due to high leaf-to-soil transfer rates. Given that seedlings emerge from the soil, it is reasonable to expect that some of the first bacterial colonists would come from soil, but that leaf-adapted microbial propagules may ultimately outcompete them. We see support for generally high similarities of corn and soybean phyllosphere microbiomes to soil bacterial communities in our data (Fig. 3B,D), but our design allows us to examine how this relationship changes over space (distance from the woodland edge as well as distance from the soil), time, host species, and leaf age. First, we see that for both corn and soybean microbiomes, the microbiomes from plants farther into the field interior are more similar to soil microbiomes (Fig 3A,C). In other words, the plants closer to vegetation sources of microorganisms tended to exhibit less similarity to soil microbiomes. This suggests that if a seedling were to emerge into a robustly colonized plant community, its initial microbiome may be more plant- rather than soil-affiliated. Second, we see that for corn plants, phyllosphere similarity to soil is lower at the final timepoint relative to the other timepoints (Fig. 3B). There could be at least two drivers underlying this effect: 1) the older leaves have had more opportunity to filter their microbiome and/or be colonized by leaf-adapted taxa, and 2) although we consistently sampled the same cohort of leaves through time, the corn plants were taller, and hence farther from the soil surface later on. Notably this change in similarity to soil was not observed in the soybean plants, which have a shorter, bushier growth habit than corn. Close spatial association of *Arabidopsis* leaves with soil was also shown to increase the likelihood of detection of soil-associated taxa with these leaves (Massoni et al., 2021). Third, we see that for both corn and soybean plants, the younger leaf microbiomes at the final sampling timepoints were more similar to those of the leaves of the same plants and conspecific neighbors than they were to the soil or the surrounding woodland vegetation (Fig. 6A,B). This observation lends further support to the idea that leaves emerging farther from the soil will share fewer taxa with soil. Lastly, we see that older corn leaves, which were closer to the soil at emergence, were more similar to soil than younger leaves, which emerged higher up on the plant (Fig. 8A), supporting the notion that distance to soil may be an important factor in soil-phyllosphere microbiome similarity. Thus the conceptual model that emerges from our set of comparisons is that leaves emerging into a habitat with a paucity of leaf-associated taxa will tend to share more microbial taxa with the soil, in part due to their close proximity to the soil, and also due to the lack of suitable colonists from vegetation, but as the surrounding plant community develops and enriches for leaf-associated taxa, leaves that emerge later will be more readily colonized by leaf-associated taxa.

Our study highlights the importance of incorporating microbial spatial ecology into agroecosystem research. While the movement of pests and pathogens in agricultural zones has long been a primary research focus (Hudelson et al., 1989; Morris and Sands, 2017; Sessitsch et al., 2023; Upper et al., 2003), there has comparably less work centered on the dispersal of beneficial or commensal microbial taxa in an agricultural setting (Grady et al., 2019; Meyer et al., 2022; Whitaker et al., 2023, 2021). Dispersal limitation of leaf-associated biota has been observed in a variety of ecosystems, including forests and orchards (Finkel et al., 2012, 2011; Galès et al., 2014; Laforest-Lapointe et al., 2016; Lajoie and Kembel, 2021; Lindow and Andersen, 1996; Lindow and Suslow, 2003; Noble et al., 2025, 2020; Yang et al., 2023), and we add to this body of work by showing that neighboring vegetation from mixed woodland/herbaceous plant communities can expel sufficient fluxes of microorganisms to effect a change in the composition, abundance, and host filtering of corn and soybean phyllosphere microbiomes grown under modern agricultural contexts that are typified by large fields devoid of other vegetation at the time of crop plant emergence. While testing the health or yield outcomes of plants along spatial gradients into these fields was beyond the scope of our study, other studies have shown that 1) crop phyllosphere colonization can be slow following emergence (Ercolani, 1991; Jacques et al., 1995; Lindow, 1982; Lindow et al., 1983, 1978; Miller et al., 2019; Thompson et al., 1993) and can proceed more rapidly if suitable microorganisms arrive on the plant (Lindow and Suslow, 2003), and 2) early and rapid colonization of the phyllosphere can protect the plant through competitive exclusion of pathogens and/or priming of plant immunity (Lindow, 1982; Lindow et al., 1983; Lindow and Suslow, 2003; Ritpitakphong et al., 2016; Vogel et al., 2016, 2021). Thus, given that many crop pathogens are found in crop residues in the soil (Cook et al., 1978; Kerdraon et al., 2019), tipping the balance of colonization towards plant- rather than soil-originating microorganisms by providing sources of vegetation-affiliated microorganisms may prove beneficial for the plant and provide co-benefits for the surrounding environment (Amy et al., 2015; Garratt et al., 2017; Staley et al., 2023).

## Acknowledgements

We thank L. Kinkel and her research group, especially A. Hughes and J. Ahlborn, for their hospitality and help at the University of Minnesota during field collection and sample processing. We thank B. Webster at the University of Minnesota Rosemount Research and Outreach Center for his assistance and expertise in identifying appropriate sites as well as providing logistical support for field collections and site meteorological data. We thank D. and N. Meyer for room, board, and access to transportation during field collection and sample processing. Lastly, we thank B. Koskella for access to lab space and key equipment for ddPCR and DNA extraction.

