## Supplementary Materials for "Microbial dispersal from surrounding vegetation influences phyllosphere microbiome assembly of corn and soybean"

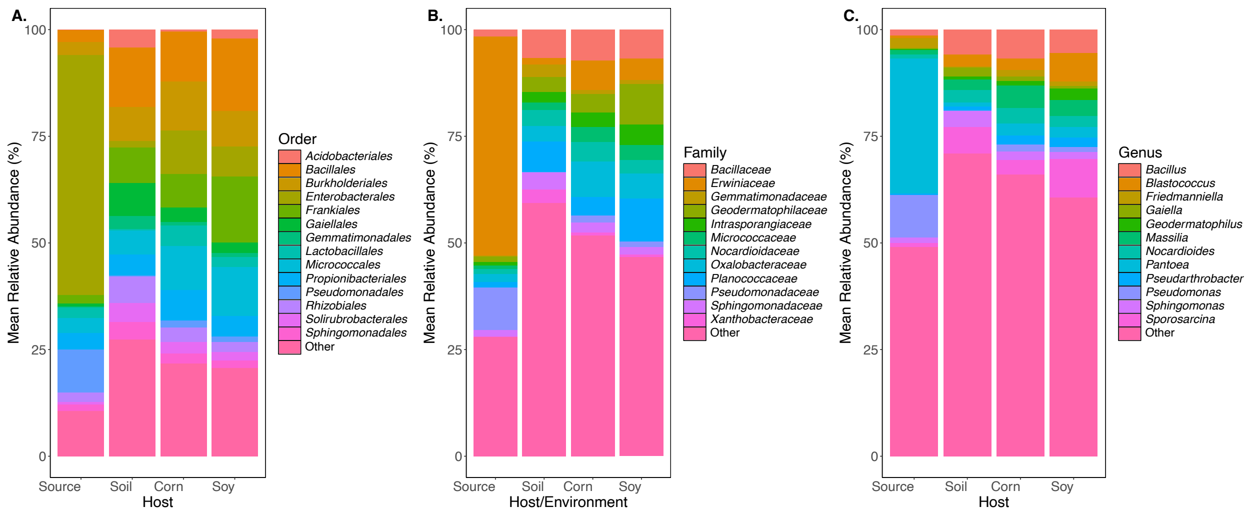


**Supp. Fig. 1:** Stacked bar chart representing the composition of the epiphytic bacterial community composition (relative abundances, y-axis) of surrounding vegetation (Source), soil, corn, and soybean plants (x-axis). Taxa with fewer than 10 occurrences have been removed. Taxonomic structure is organized at the A) order level, B) family level, and C) genus level. Any orders or families with less than 3% relative abundance were labelled as Other, and any genera with less than 2% relative abundance were labelled as Other.

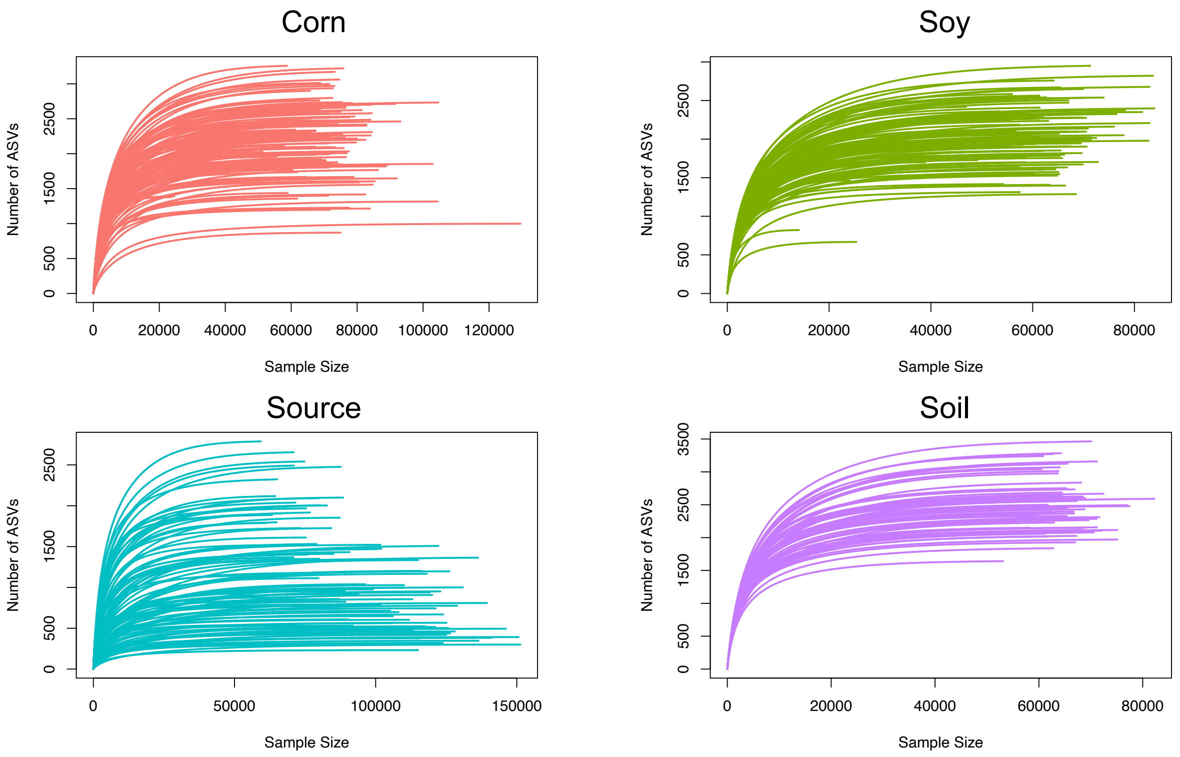


**Supp. Fig. 2**: Rarefaction curves for corn, soy, source vegetation, and soil bacterial microbiomes demonstrating the relationship between sampling depth (x-axis) and number of bacterial ASVs detected (y-axis). Each line represents one sample. Note that x-axis differs among plots.

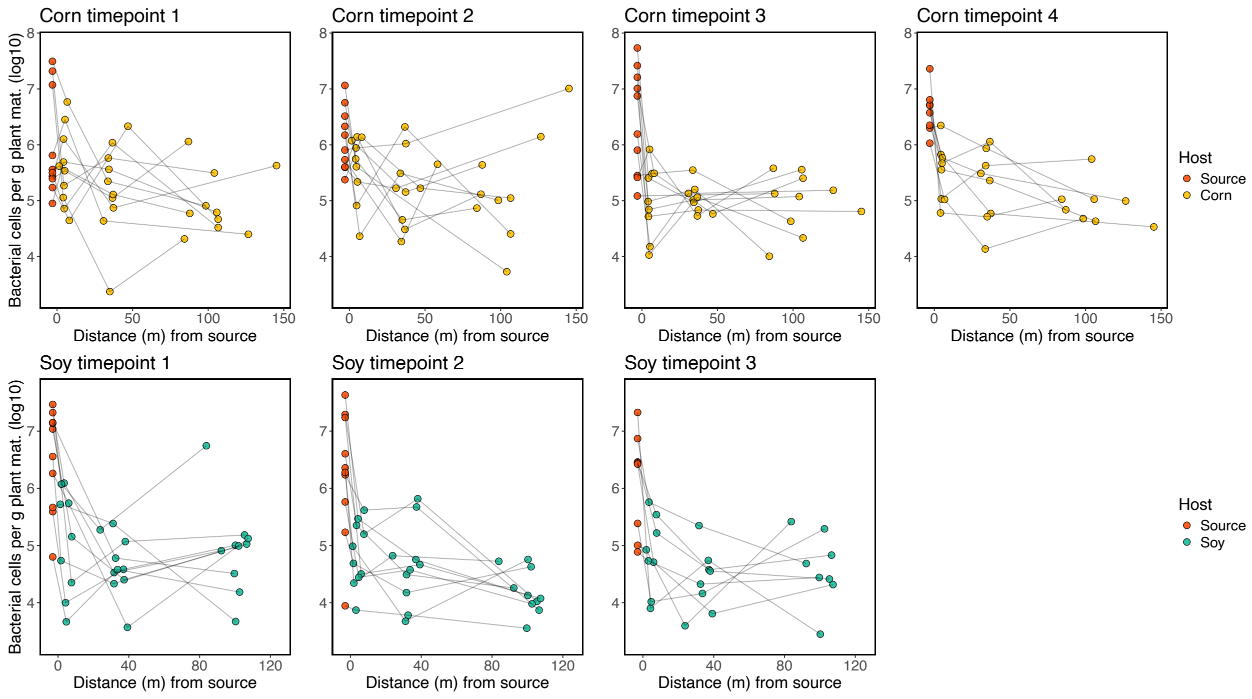

**Supp Fig 3:** Bacterial cell densities of epiphytic microbiomes decline over distance from the source vegetation surrounding fields (red points). Lines connect samples of the same transect.

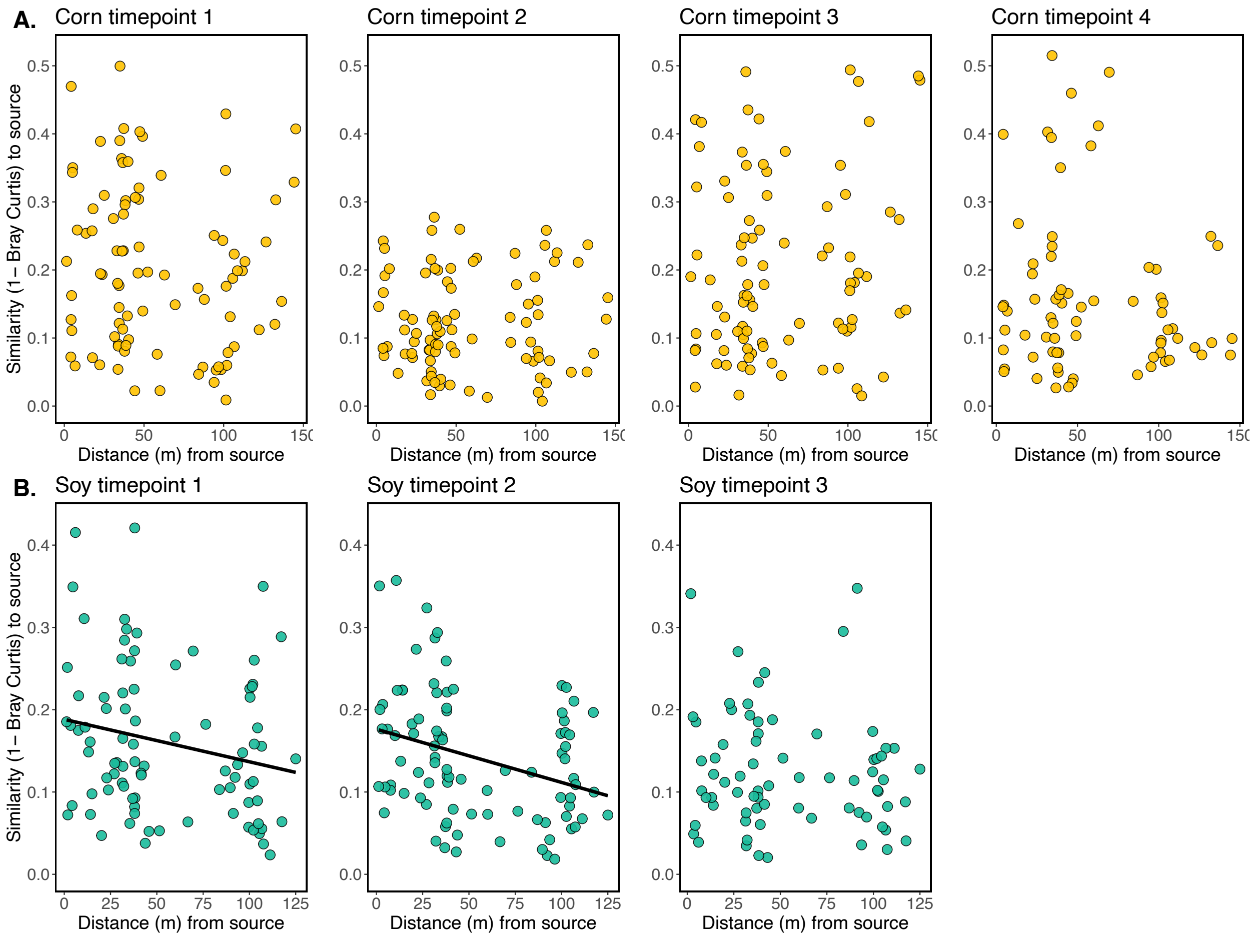


**Supp. Fig 4**: The relationship between epiphytic microbiome similarity to surrounding vegetation (1-Bray Curtis, y-axis) and distance (m) from that vegetation (x-axis). No soil-affiliated taxa were removed from these data. This supplementary figure corresponds to Fig. 4, in which soil-affiliated taxa were removed. Solid fit lines are shown where a statistically significant ($p < 0.05$) relationship is observed.

### Absolute abundance of vegetation–affiliation bacteria

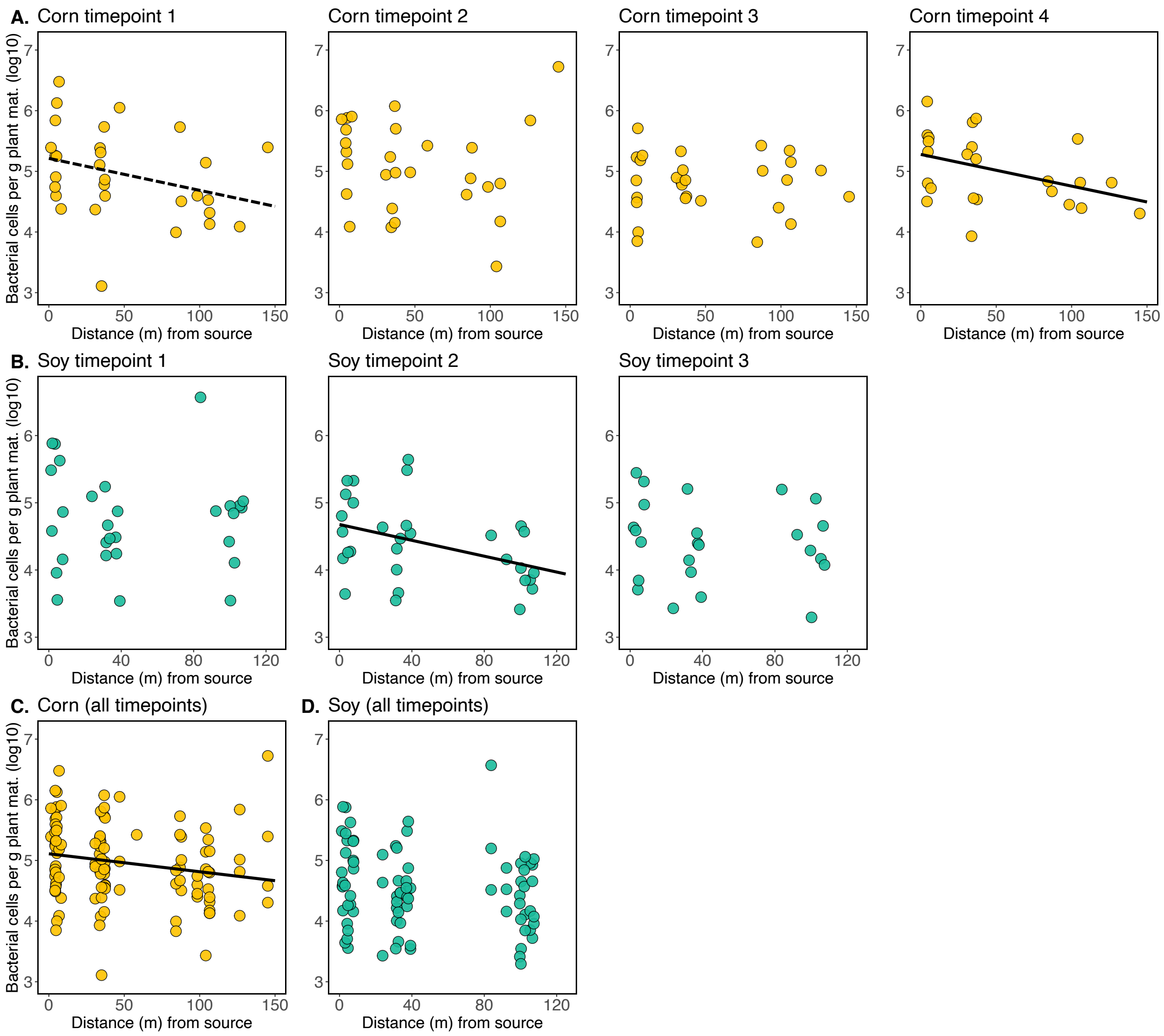


**Supp Fig. 5**: The estimated absolute abundance of surrounding vegetation-affiliated bacteria (y-axis) over distance (x-axis). A) Corn samples over timepoints 1-4. B) Soybean samples over timepoints 1-3. C) Corn samples with all timepoints combined. D) Soybean samples with all time points combined. Solid lines indicate statistically significant ($p$ <0.05) linear model fit in which cells per gram density is the response variable, geographic distance is the fixed effect, and site is a random effect. The dashed line indicates a statistically trending ($p$ = 0.07) relationship.

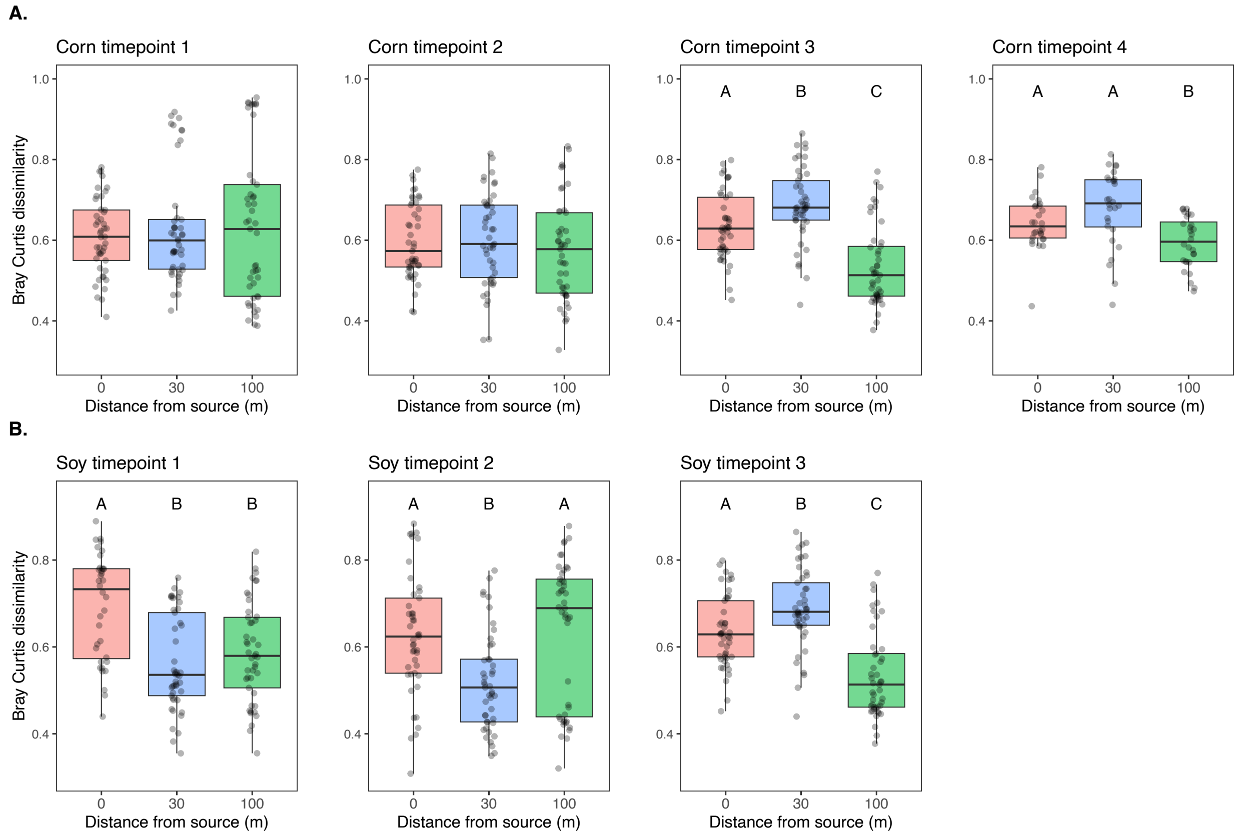


**Supp Fig. 6:** The relationship between Beta diversity (Bray Curtis dissimilarity, y-axis) and distance from the surrounding (source) vegetation (x-axis) for corn (A) and soybean (B) epiphytic bacterial communities. Plots are separated by time points. Letters above boxes indicate statistically significant differences among groups ($p < 0.05$, Tukey's HSD).

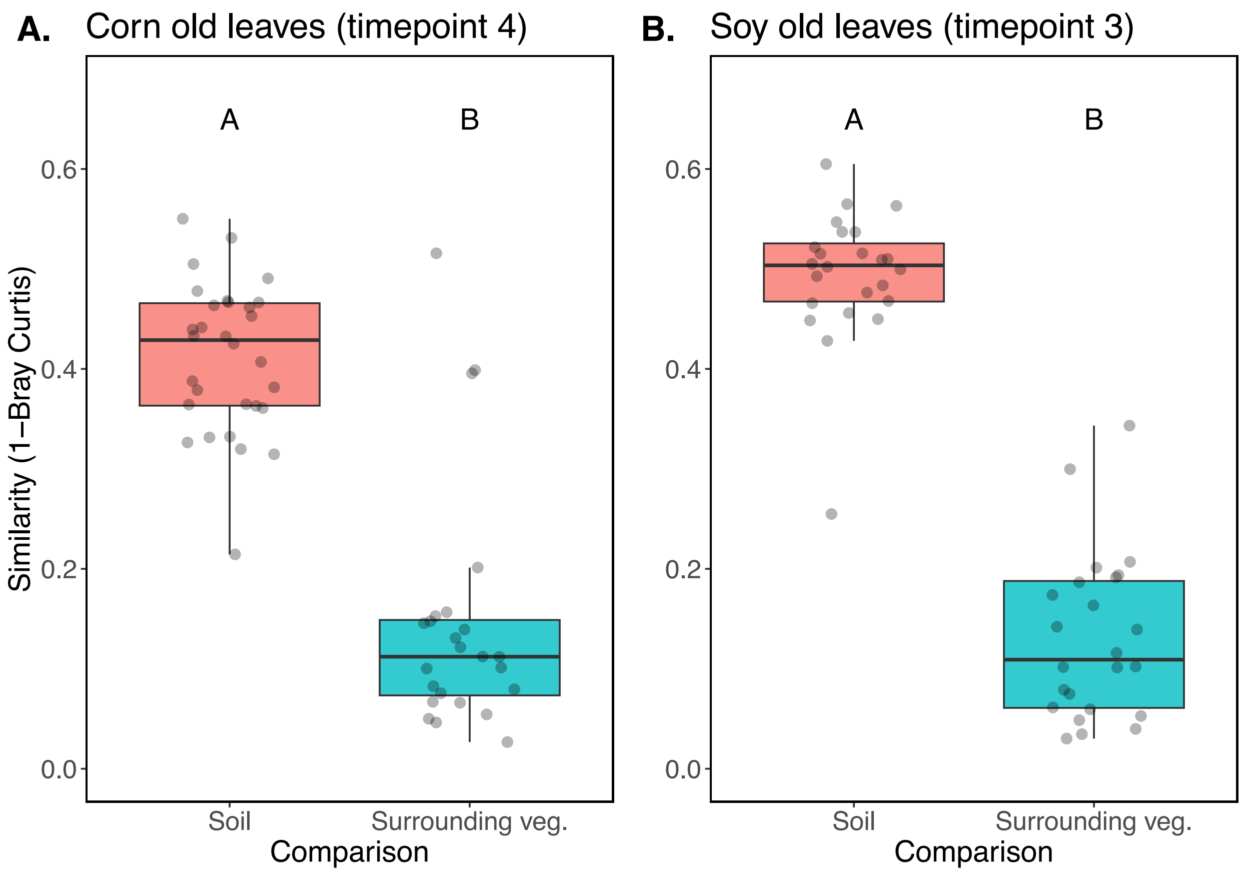


**Supp Fig. 7**: Taxonomic similarity (1-Bray Curtis, y-axis) of epiphytic microbiomes from the last timepoint of old corn leaves (lowest on the plant) (left plot) and old soybean leaves (lowest on the plant) (right plot) to two putative sources of microbial propagules (x-axis): soil from the paired sampling locations (Soil) and leaves from the vegetation surrounding the field sampled at the same transect (Surrounding veg.).

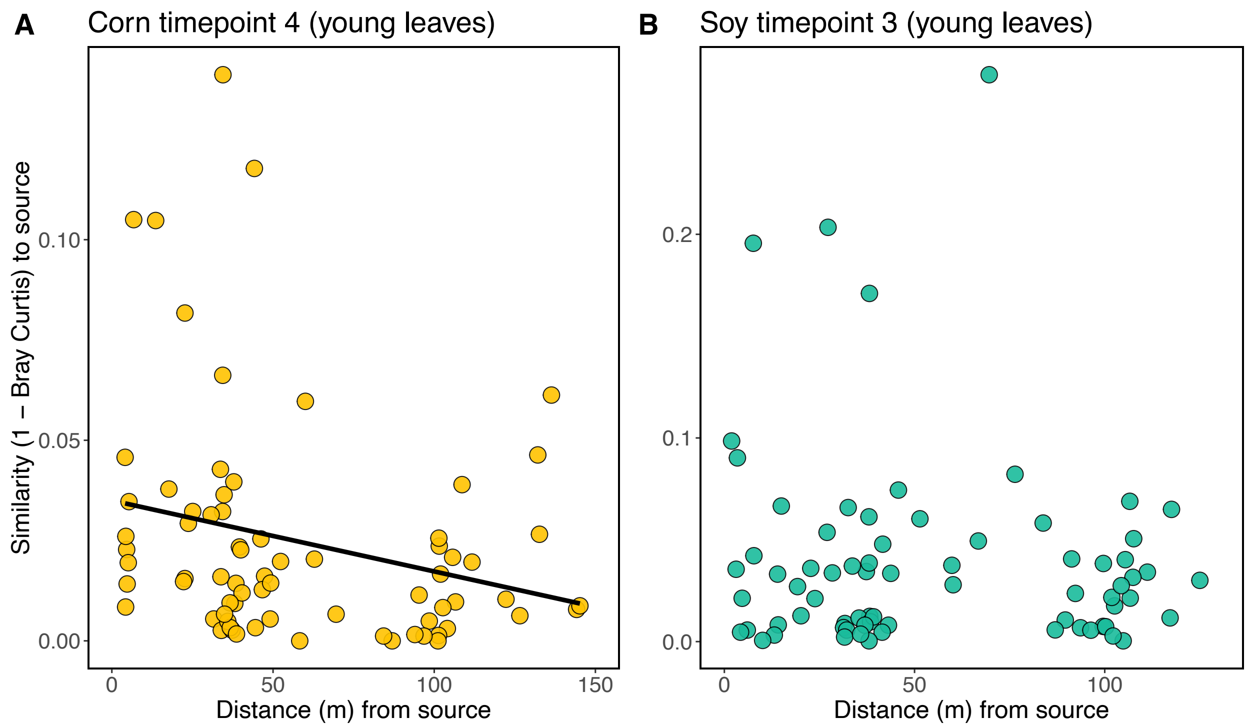


**Supp. Fig. 8**: Relationship between epiphytic microbiome similarity of young leaves to surrounding vegetation (1-Bray Curtis, y-axis) and distance (m) from that vegetation (x-axis). Soil-affiliated taxa were removed from these data. The solid fit line is shown where a statistically significant ($p$ < 0.05) relationship is observed.

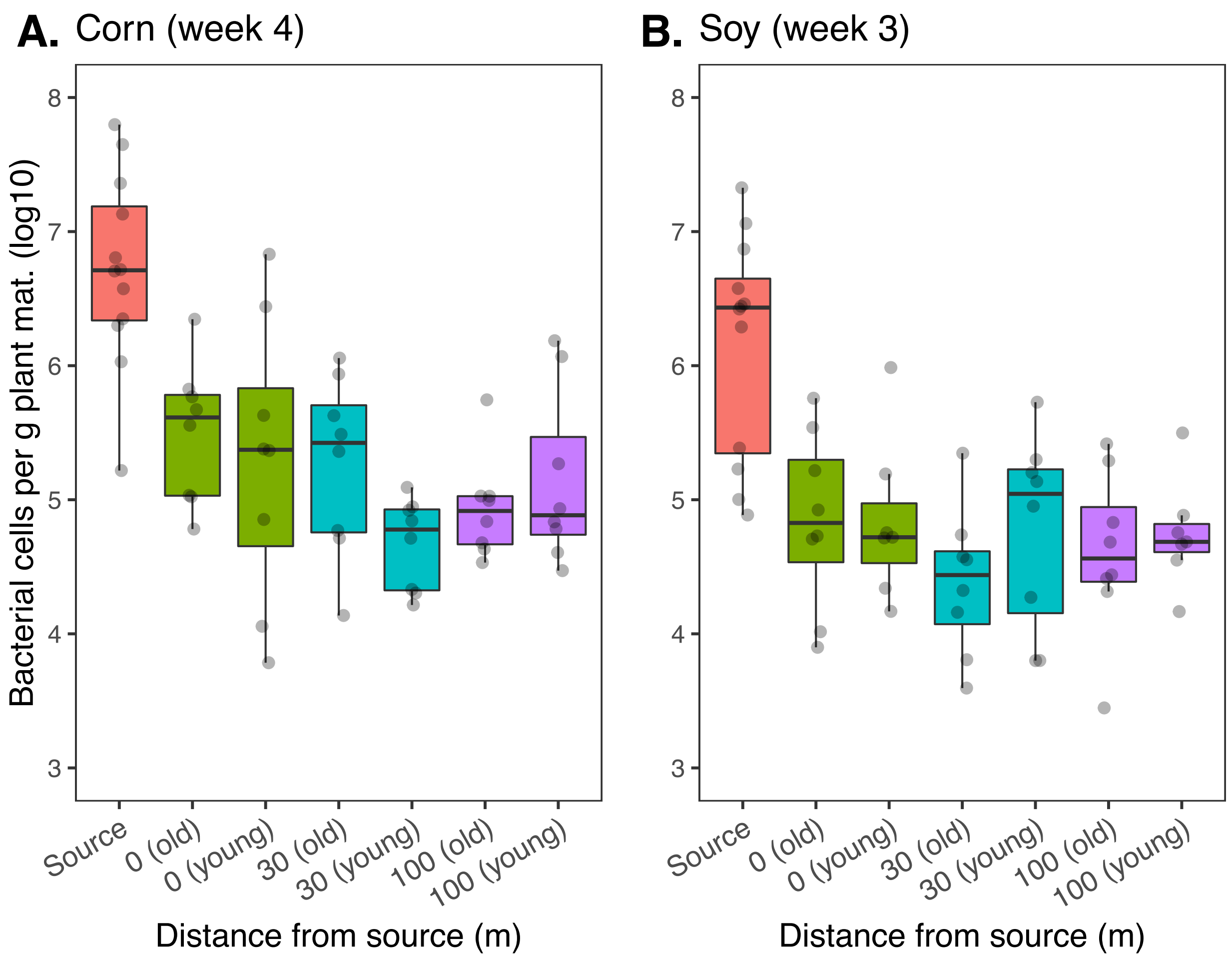


**Supp Fig. 9**: Bacterial cells per gram of plant material (y-axis) on young and old leaves over distance from the surrounding vegetation (x-axis) for corn (A) and soybean (B) samples taken at the last sampling timepoint.
